# A Reference Landscape of Regulatory T Cell States in Mice

**DOI:** 10.64898/2026.01.30.702856

**Authors:** A Freuchet, N. Mehrotra, I. Magill, O. Casey, X. Chi, S. Zhang, C.J. Imianowski, J.A. Myers, J.R. Huh, L. Brossay, D.A.A. Vignali, C Benoist, D. Zemmour, the immgenT project

## Abstract

CD4⁺FoxP3⁺ regulatory T cells (Tregs) are central to immunity, tolerance, and tissue homeostasis, yet their extensive heterogeneity lacks a unifying framework. Within the immgenT project, we profiled gene expression, surface markers and TCR clonotypes of mouse Tregs. Using a joint RNA–protein deep generative model, we define the Treg landscape, organized around eight conserved clusters shared across tissues and conditions, with immune context reshaping their relative abundance rather than generating new states, including a prominent circulating effector Treg population enriched in select non-lymphoid tissues. We validate this framework by integrating external datasets from conditions not represented in immgenT and by defining a flow cytometry panel spanning the Treg landscape. Together, immgenT provides a scalable, reusable reference that unifies Treg heterogeneity across tissues and immune challenges.

## Introduction

CD4^+^ FoxP3^+^ regulatory T cells (Tregs) are essential for enforcing immune tolerance and maintaining tissue homeostasis. Their biology extends far beyond classical immunosuppression, encompassing metabolic regulation, control of inflammation, and tissue repair1. This functional breadth is reflected in substantial heterogeneity across Tregs, encompassing differences in transcriptional programs, surface marker expression, developmental origin, antigen specificity, and tissue localization, among other features^1–8^.

Multiple layers of Treg heterogeneity have been described. Ontogenetically, thymus-derived Tregs (tTregs) and peripherally induced Tregs (pTregs) arise through distinct developmental pathways and contribute differentially to immune tolerance, particularly at barrier sites^9–14^. Neonatal Tregs represent an additional wave of regulatory cells with unique properties that influence lifelong immune balance^15^. Spatial context further shapes Treg identity: Tregs residing in non-lymphoid tissues (NLTs), including visceral adipose tissue (VAT), skeletal muscle, colon, skin, lung, liver, kidney, central nervous system, and reproductive organs, exhibit specialized transcriptional signatures, distinct T cell receptor (TCR) repertoires, and tissue-adapted functions^1,4,7^.

Single-cell RNA sequencing has been instrumental in uncovering this diversity, revealing classical resting-effector axes as well as multiple effector Treg states^6,9,16–22^. In many contexts, Tregs acquire transcription factor programs that mirror those of the effector T cells they suppress or reflect the inflammatory environments in which they reside. For example, expression of T-bet and CXCR3 equips Tregs to restrain Th1 responses (i.e., Th1-Tregs)^18,23,24^, Bcl6 defines T follicular regulatory cells (Tfr)^25^ that control germinal center reactions, and RORγt expression in Tregs at barrier tissues shapes tolerance to commensal microbiota^11,16^.

In addition to transcriptional specialization, Tregs deploy a wide range of effector and suppressive mechanisms. While molecules such as CTLA-4 and IL2RA are broadly expressed as part of the core Treg identity^2,3,26,27^, the deployment of downstream effector pathways varies across tissues and states, encompassing suppressive cytokines (e.g., IL-10, TGF-β, IL-35), metabolic regulation, cytolytic mechanisms, and tissue-repair mediators such as amphiregulin^1,8^.

Despite this progress, the full extent of Treg heterogeneity remains incompletely defined, and the field lacks a unified framework that integrates Treg diversity across tissues and immune contexts. As most studies focus on individual organs or conditions, it remains difficult to reconcile findings and to determine whether newly described Treg populations represent distinct cell states or context-dependent manifestations of shared regulatory programs.

The immgenT project^28^ generated a comprehensive single-cell atlas of mouse T cells. This resource profiled nearly 700,000 T cells across hundreds of lymphoid and non-lymphoid tissues, homeostatic and inflammatory conditions, and diverse immune perturbations, using single-cell RNA sequencing coupled with paired CITE-seq and TCR sequencing. Within this atlas, more than 44,000 Tregs are captured, enabling systematic resolution of Treg diversity at unprecedented scale. We show that Treg heterogeneity can be organized into a remarkably simple and robust framework comprising eight conserved clusters that span a continuous state space. Together, these clusters define a shared repertoire of Treg states whose relative abundance varies across tissues and inflammatory contexts, indicating that context primarily modulates the balance of conserved states rather than generating entirely new ones. By mapping external datasets using reference-based integration (T-RBI, a custom algorithm introduced in detail in the accompanying immgenT-Cosmology manuscript^29^) and by defining an eight-marker flow cytometry panel that recapitulates atlas-defined states, immgenT provides a reusable reference framework and common language for the study of Treg biology across experimental systems.

## Results

### Scope and overview of Treg heterogeneity in immgenT

As part of the immgenT effort to comprehensively chart mouse T cell states, we sequenced 44,380 Treg cells within a total of 682,951 CD3⁺ T cells collected across 658 samples. Although baseline secondary lymphoid organs (SLOs) from unmanipulated WT B6 mice (6–8 weeks, both sexes) contributed a substantial fraction of Tregs, a large proportion originated from non-lymphoid tissues (39%) or from immune perturbation settings (56%), spanning models of infection, autoimmunity, immunization, cancer, and physiological states such as pregnancy and aging **(Extended Data Table 1; Extended Data Fig.1a-c**).

As described in the companion immgenT-Cosmology manuscript^29^, all T cells were integrated using totalVI, a state-of-the-art deep learning framework that jointly models RNA and protein measurements to define a shared latent space and projected into a unified embedding of all T cells (the all-T Minimum Distortion Embedding; MDE). Within this global T cell landscape, Treg cells formed a distinct population, clearly separated from conventional CD4⁺ T (Tconv) cells and positioned closer to activated Tconv. This positioning reflects the basal activation state of Treg cells, in contrast to the quiescent phenotype of resting Tconv (**Fig. 1a**).

**Figure 1.**
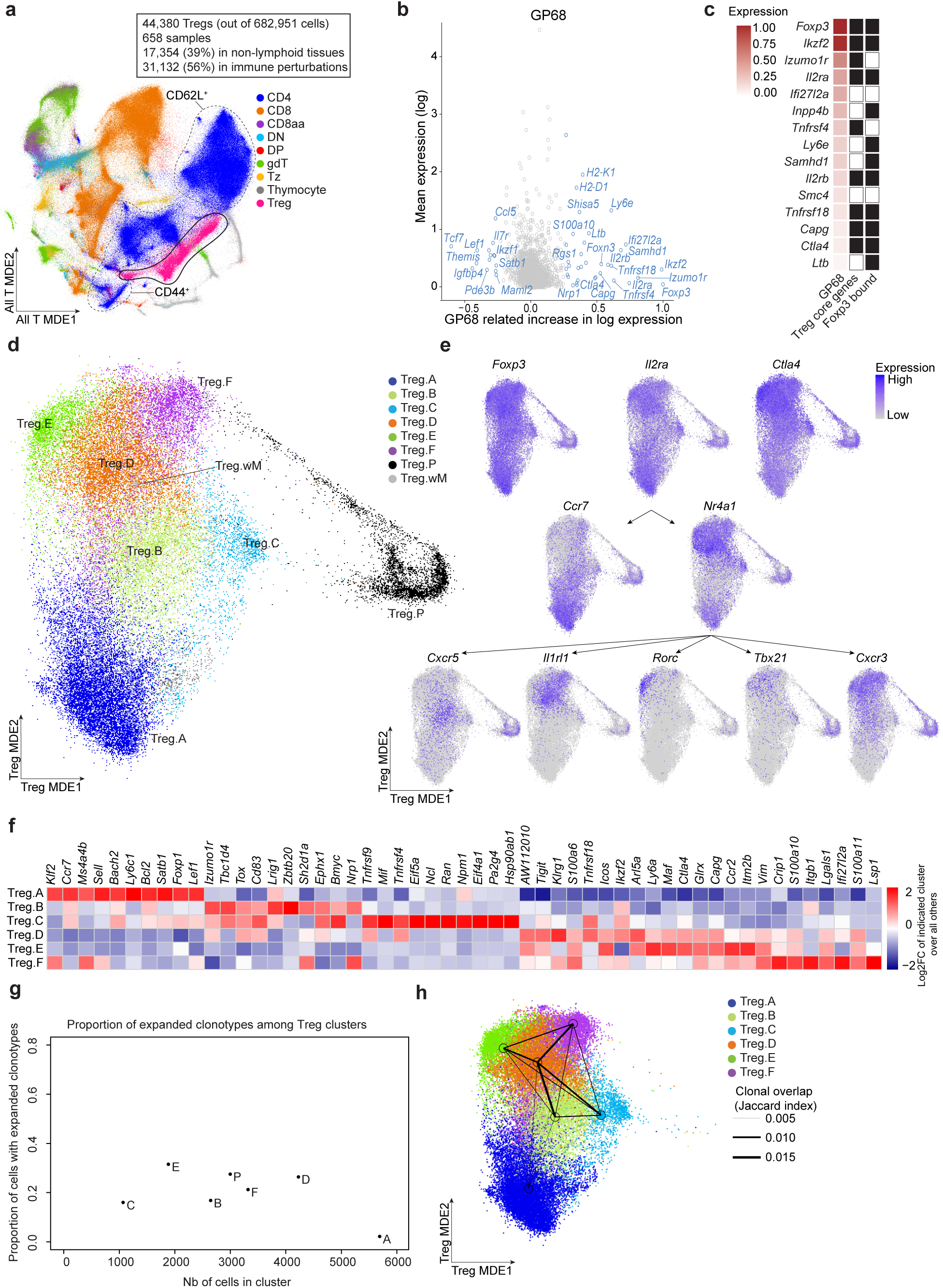
immgenT resolves Treg heterogeneity into eight stable clusters. **(a)** Minimum Distortion Embedding (MDE) of all T cells in immgenT (all-T MDE), colored by major T-cell lineages (CD4, CD8, CD8aa, DN, DP, gd T cells, Tz (Zbtb16^+^) cells, and Treg). A total of 682,951 T cells, including 44,380 Tregs, were profiled across diverse tissues and immune conditions using single-cell RNA sequencing with paired CITE-seq (128-plex) and αβ TCR sequencing. **(b)** Mean expression versus fold-change plot for GP68, the Treg-specific Gene Program. Values are log-normalized, and fold changes are scaled relative to the maximal induction. **(c)** Heatmap showing the expression of the top 15 genes in GP68. Genes are annotated for inclusion (black) or absence (white) in previously defined core Treg genes^21^ and Foxp3 ChIP-seq targets^30^. **(d)** Treg-specific MDE (Treg-MDE) of all Tregs, colored by cluster assignment. **(e)** Treg-MDE displaying expression of genes commonly associated with Treg heterogeneity, including *Foxp3, Il2ra, Ctla4, Ccr7, Nr4a1, Cxcr5, Il1rl1, Rorc, Tbx21, and Cxcr3*. **(f)** Heatmap showing expression of the top 10 genes per cluster, ranked by log2 fold change relative to all other Treg clusters. **(g)** Scatterplot of the proportion of cells with expanded clonotypes versus cluster size. **(h)** Graph representation of clonotype sharing overlaid on the Treg MDE. Nodes are positioned at cluster centroids; edge width is proportional to the extent of clonotype sharing (Jaccard index).

An independent analysis based on gene-program (GP) discovery using empirical-Bayes matrix factorization (immgenT-Cosmology manuscript^29^) (GP can be interpreted as a reference-free gene signature) likewise identified Treg cells as a coherent population through a single, highly specific program, GP68, present in virtually all Treg cells and rarely expressed in other T cell states (**Extended Data Fig. 1d**). GP68 encompassed canonical Treg markers, including *Foxp3, Il2ra, Izumo1r, Tnfrsf18* (GITR), and *Ctla4* (**Fig. 1b**). These genes substantially overlapped with previously defined Treg core genes^21^ and Foxp3-bound loci identified by ChIP-seq in *Foxp3*-deficient mice^30^ (**Fig. 1c**).

To resolve finer-grained structure within the Treg compartment, we generated a Treg-specific dimensionality reduction plot, providing a standardized visualization of Treg states ("Treg MDE") (**Fig. 1d**). This representation revealed a largely continuous Treg state space, reflecting the broad diversity of tissues, immune challenges, and experimental conditions captured in the dataset. Despite this continuity, clustering performed jointly in transcriptomic and protein space robustly identified six major Treg clusters, along with a proliferative cluster (**Extended Data Fig.1e**) and a small rare-state “miniverse” cluster. The “miniverse/wM” state (1.7% of Tregs) seems to correspond to a shared stressed transcriptional state observed across multiple T cell lineages; its biological significance remains unclear and is discussed in the immgenT-Cosmology manuscript^29^.

Well-characterized Treg populations were readily recovered with expected transcriptional markers (**Fig. 1e,f, Extended Data Table 2 with cluster gene signatures**), and characteristic tissue enrichments (**Extended Data Fig. 1f**). Cluster A represented resting Tregs (rTregs), enriched in spleen and marked by high expression of *Sell, Bach2, Ccr7, Satb1,* and *Lef1*. Effector Tregs (*Nr4a1*⁺) were heterogeneous. Cluster D comprised *Il1rl1* (ST2)⁺ *Klrg1*⁺ Tregs enriched in NLTs and previously linked to tissue homeostasis and repair^31,32^, whereas cluster E expressed *Rorc* and included gut-associated Tregs involved in microbiota tolerance^11,12,33^. Cluster B contained T follicular regulatory (Tfr) cells, expressing *Cxcr5* and *Pdcd1,* and was enriched in lymph nodes during NP–OVA immunization. Cluster C exhibited a strong TCR-response signature with fewer effector features than clusters D, E, or F, consistent with observations in another dataset^9^ (**Extended Data Fig. 1g**). Cluster F was characterized by expression of *Itgb1* (CD29), *Itga2* (CD49b), and *Klf2*, representing a recurrent Treg state discussed further in Figure 6. Notably, T-bet⁺ Tregs did not form a discrete cluster; instead, *Tbx21* and *Cxcr3* expressions were distributed across multiple effector subsets (**Fig. 1e**). Effector molecules, while most strongly expressed in activated clusters, remained broadly distributed across multiple states (**Extended Data Fig. 1h**). Activated clusters were the most clonally expanded populations, with approximately 20% of cells in each cluster belonging to expanded clonotypes, and they exhibited substantial clonotype sharing, consistent with shared ancestry followed by heterogeneous differentiation or plasticity (**Fig. 1g,h**).

pTregs did not form separate clusters. TCR clones shared between Tconv and Tregs, indicating shared ancestry and compatible with pTregs or ex-Tregs, were distributed across all Treg clusters, with some depletion in Treg.A and enrichment in Treg.E and Treg.P compared with Treg-specific clones from the same experiments (**Extended Data Fig. 2a,b**). Using T-RBI, we also mapped a published dataset^34^ of bona fide pTregs differentiated *in vivo* from Foxp3-negative Tconv cells, and these pTregs likewise adopted a broad range of transcriptional phenotypes (**Extended Data Fig. 2c**).

Overall, immgenT provides a unified framework for resolving Treg heterogeneity across tissues and immune perturbations. The substantial sharing of clonotypes, suppressive programs, and tissue-adaptive features across activated clusters argues against strict context-dependent specialization of individual Treg states.

### Bridging flow cytometry and single-cell RNAseq: An 8-marker surface panel resolves Treg heterogeneity

Leveraging the 128-antibody CITE-seq panel (full panel description in Extended Data Table 2 from the immgenT-Cosmology paper^29^), we revisited surface marker expression patterns distinguishing Treg cells from conventional CD4⁺ T cells and among Treg clusters (**Fig. 2a**). Consistent with classic work, Treg cells expressed higher levels of CD25, GITR, FR4, CD73, and Neuropilin-1 (NRP1), and lower levels of CD45RB compared to Tconvs^2,35^ (**Fig. 2a**). However, none of these markers were uniformly expressed across all Tregs. CD25^hi^ IL-7Rα^lo^ gating preferentially captured resting Tregs and excluded many effector populations (**Extended Data Fig. 3a**), consistent with IL2RA downregulation upon activation. Conversely, CD25^lo^ IL7Rα⁺ Tregs were enriched for activated and effector states (**Extended Data Fig. 3b**). CD45RB^hi^ gating also more effectively enriches resting Tregs than the commonly used CD45RB^lo^ gate (**Extended Data Fig. 3c,d**).

**Figure 2.**
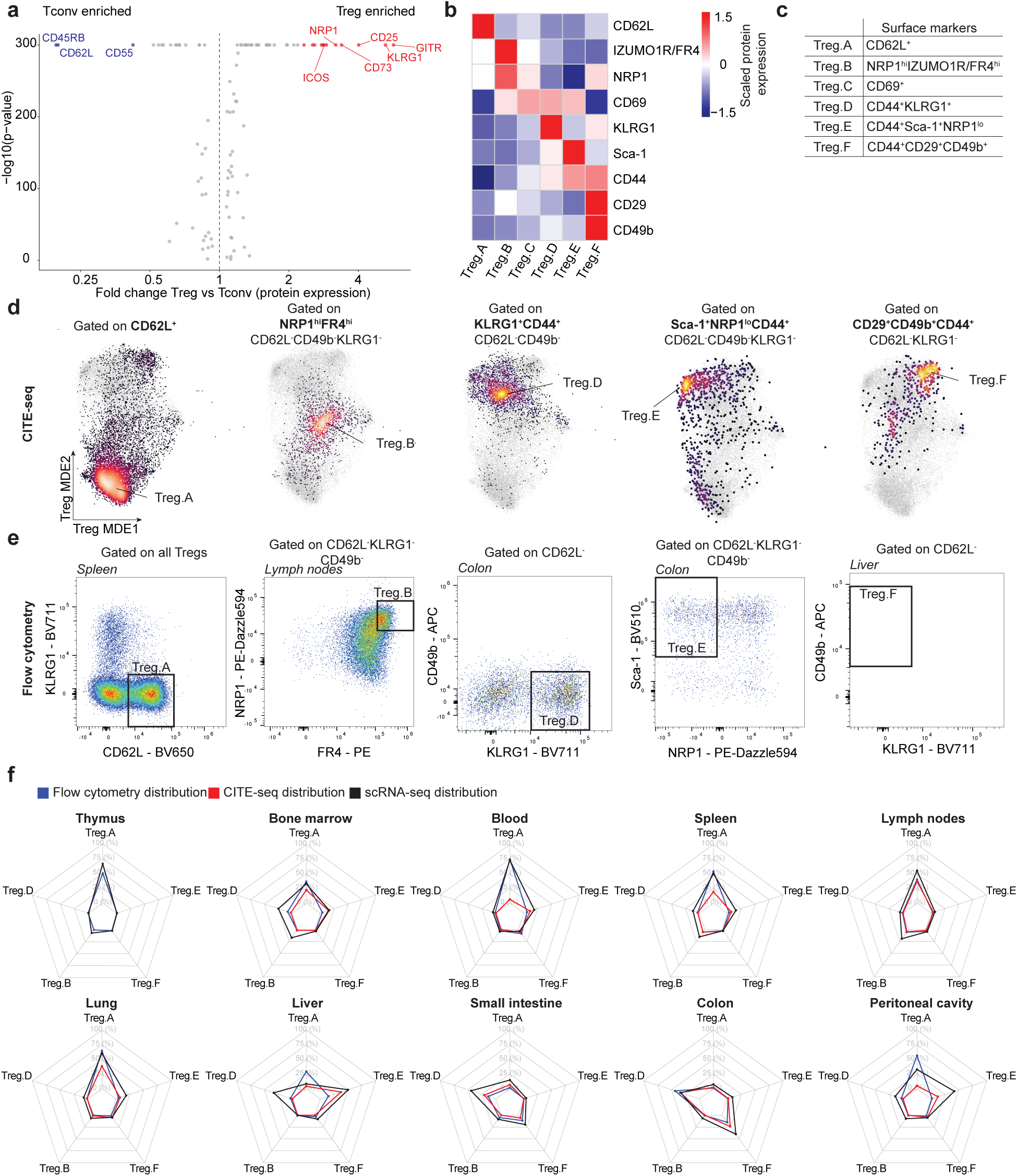
An eight-color flow cytometry panel resolves Treg clusters. **(a)** Volcano plot showing differential protein expression by CITE-seq between Tregs and conventional CD4⁺ T cells, plotted as log2 fold change versus -log10(*P* value). (b) Heatmap showing the expression of the most discriminative surface markers across Treg clusters, measured by CITE-seq (Z-score). **(c)** Summary table of gating strategies used to identify individual Treg clusters based on surface markers. **(d)** Treg-MDE highlighting cells bioinformatically gated using the CITE-seq strategies described in (c). **(e)** Representative flow cytometry plots from spleen, lymph nodes, colon, and liver showing the same gating strategies as in (c). Data are representative of 3–4 independent experiments. **(f)** Radar plots comparing the proportion of each Treg cluster identified by scRNAseq (black), CITE-seq (red), and by flow cytometry (blue) across organs. Values are shown as median ± SEM (CITE-seq: n = 2–55 samples per organ; flow cytometry: n = 3–5 experiments per organ).

We next asked whether a flow cytometry–style gating strategy could be adapted to resolve the Treg heterogeneity revealed by high-dimensional single-cell RNA-seq. Through a systematic analysis of the sensitivity and specificity of all 128 surface markers for distinguishing Treg clusters (see **Methods**), we defined a concise 8-marker panel, including CD62L, CD44, CD69, KLRG1, NRP1, IZUMO1R/FR4, CD49b, and Sca-1 (**Fig. 2b,c** **and Extended Data Table 3**). Cluster A corresponded to CD62L⁺ CD44^low^ resting Tregs. CD62L⁻ CD44⁺ effector Tregs encompassed clusters D, B, E and F, which could be further distinguished as follows: Treg.D (KLRG1⁺ CD49b⁻), Treg.E (Sca-1⁺ NRP1^lo^), and Treg.F (KLRG1⁻ CD49b⁺). For Treg.B, corresponding to T follicular regulatory cells, NRP1^hi^ FR4^hi^ gating provided an effective alternative to the classical CXCR5/PD- 1 combination. Treg.C was best defined as CD62L⁻ CD69⁺ Tregs lacking KLRG1, CD49b, and Sca-1. Applying these gating strategies directly to the CITE-seq data—as would be done in conventional flow cytometry—consistently highlighted the expected regions of the Treg MDE (**Fig. 2d**), demonstrating that this surface marker logic specifically recovers Treg states defined by high-dimensional single-cell RNA-seq.

We next validated this 8-marker panel by flow cytometry across 10 tissues (spleen, lymph nodes, blood, bone marrow, thymus, liver, colon, small intestine, and peritoneal cavity). Marker discrimination was generally stronger by flow cytometry (**Fig. 2e,f, Extended Data Fig. 4**), with improved signal-to-noise ratios enabling clearer separation of positive/negative and high/low populations and increased cluster purity (**Extended Data Fig. 5a**). Across tissues, cluster distributions quantified by flow cytometry closely matched those obtained by scRNAseq and CITE-seq (**Fig. 2f; Extended Data Fig. 5b**). As expected, no surface marker specifically identified proliferating Tregs; instead, these cells were reliably detected using intracellular Ki67 staining (**Extended Data Fig. 5c**), consistent with the specificity of *Mki67* transcripts (**Extended Data Fig. 1e**). We further confirmed that CD62L⁻ KLRG1⁻ NRP1^lo^ Sca-1⁺ Tregs were enriched in the gut and expressed RORγt, consistent with the well-established microbiota-dependent RORγt⁺ Treg population (**Extended Data Fig. 5d,e**).

Overall, the immgenT-Treg flow cytometry panel recapitulates the major axes of Treg heterogeneity revealed by single-cell RNA-seq. Through the immgenT web portal Rosetta (rosetta.immgen.org), users can go beyond this 8-marker panel to interactively explore the CITE-seq data, apply custom gating strategies that reflect their own flow cytometry experiments using any of the 128 markers, and visualize where gated populations map within the Treg MDE, positioning the Treg MDE as a shared coordinate system bridging flow cytometry and single-cell transcriptomics.

### Resting Tregs (Treg.A) are broadly distributed across tissues, including non-lymphoid organs

Resting versus effector Tregs represent a common axis of Treg biology, historically defined by gradients of CD62L/CCR7 and CD44 expression^36,37^. CD62L⁺ CD44^low^ rTregs mapped to Treg.A in immgenT (**Fig. 2d**) and was most abundant in blood (70%), lymph nodes (50%), bone marrow (39%), and spleen (47%) (**Fig. 3a**). But Treg.A was not restricted to SLOs (**Fig. 3a,b)** and constituted 1-32% of Tregs in NLTs (**Fig. 3a**). The exception was the lung, where Treg.A frequencies appeared to reach almost 60%. Most tissues were extensively perfused before processing, but residual blood contamination remained a potential explanation for the Tregs detected in NLTs. To directly test this hypothesis, we performed *in vivo* intravascular (IV) labeling by injecting anti–CD45-PE three minutes prior to sacrificing the mice, followed by *ex vivo* CD45- BUV496 staining. Blood contamination was substantial in lung and liver (>80%) but minimal in colon, lymph nodes, and spleen (<1%) (**Fig. 3c,d**). In spleen and liver—tissues with fenestrated sinusoidal capillaries—we also observed an IV-intermediate population likely representing intraparenchymal but perivascular cells. This intermediate population was not observed in lymph nodes, lung, or colon (with continuous capillaries) (**Fig. 3c,d**). These data indicate that the high apparent frequency of Treg.A in the lung arises largely from blood contamination. Nevertheless, a genuine intraparenchymal rTreg compartment persisted: 4.3% and 5.7% of extravascular lung and liver Tregs were CD62L^+^, respectively (**Fig. 3e**).

**Figure 3.**
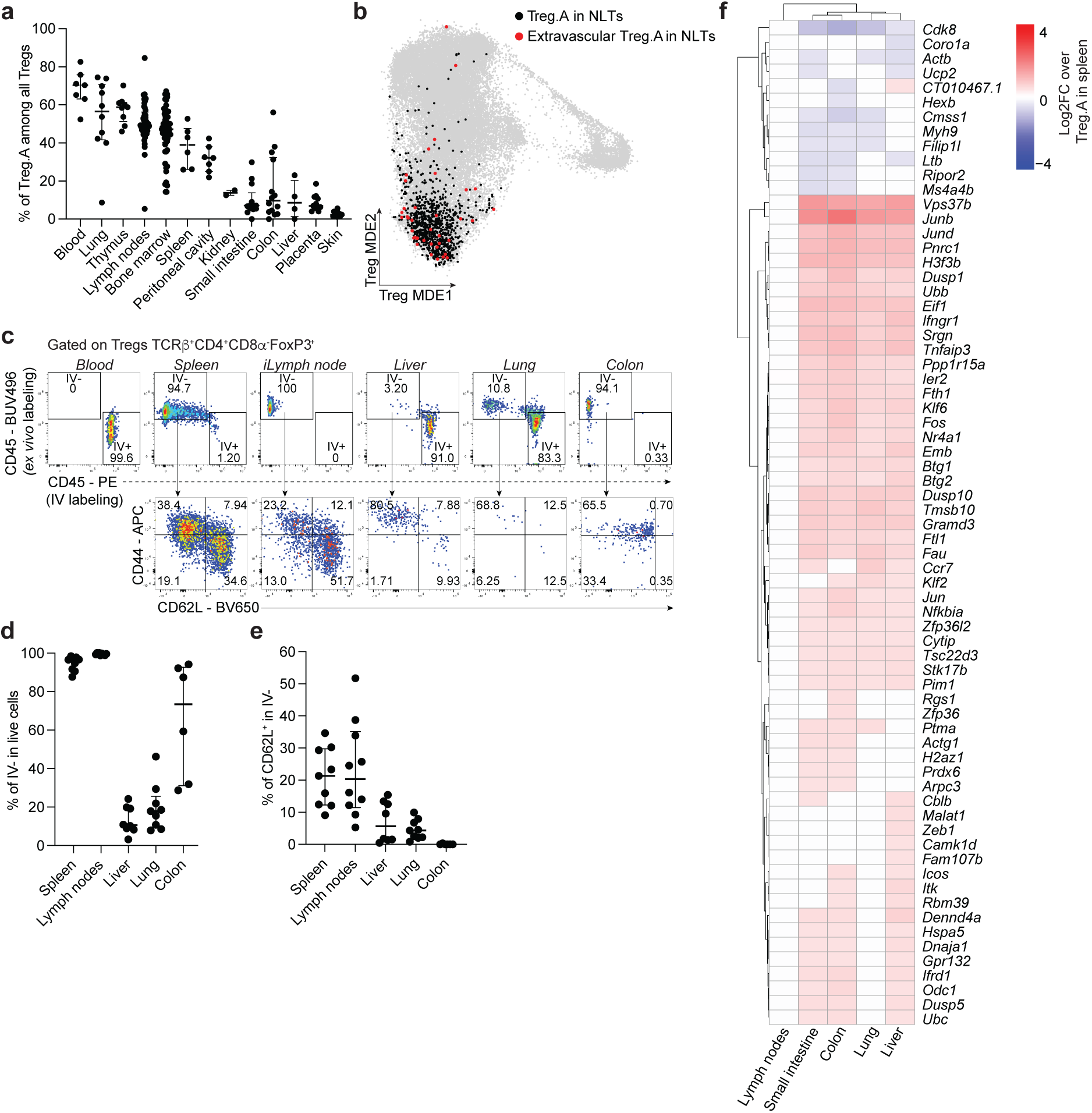
Treg.A (resting Tregs), while enriched in lymphoid tissues, are broadly distributed and present in non-lymphoid organs. **(a)** Proportion of Treg.A among all Tregs across healthy tissues in immgenT. Summary statistics show the median and interquartile range.> (b) Treg-MDE highlighting Treg.A in NLTs (colon, liver, lung, placenta, skin, and small intestine) in black. Extravascular Treg cells (IV⁻) from the same tissues are shown in red. **(c)** Intravascular versus extravascular distribution of Tregs in blood, spleen, inguinal lymph nodes, liver, lung, and colon, quantified by intravascular (IV) labeling. IV labeling was performed by intravenous injection of CD45–PE 3 min before tissue harvest (see **Methods**), followed by *ex vivo* staining with CD45–BUV496. Representative flow cytometry plots of CD45–PE versus CD45– BUV496, with gates defining intravascular (IV⁺) and extravascular (IV⁻) fractions and subsequent CD44 and CD62L expression in extravascular (IV⁻) Tregs (TCRβ⁺CD4⁺CD8α⁻FoxP3⁺) from blood, spleen, inguinal lymph node, liver, lung, and colon (n = 6–9). Data are representative of four independent experiments. **(d)** Proportion of the extravascular (IV⁻) TCRβ⁺CD4⁺CD8α⁻FoxP3⁺ Tregs across organs. Data representative of four independent experiments. **(e)** Proportion of resting CD62L^+^ cells in extravacular (IV⁻) Tregs (TCRβ⁺CD4⁺CD8α⁻FoxP3⁺) (n = 6–9). Data are representative of four independent experiments and shown as median ± SEM. **(f)** Heatmap showing differential gene expression of Treg.A across organs. Log2 fold changes are shown for lymph nodes, the small intestine, the colon, the lung, and the liver relative to the spleen. Genes shown meet the criteria of fold change >2 or <0.5 with adjusted p value <0.05 in at least one condition. *iLN, inguinal lymph node*.

We next asked whether Treg.A cells from different organs shared the same transcriptional signature (**Fig. 3f**). Compared to spleen Treg.A, lymph node Treg.A showed minimal differences . In contrast, colon, and small intestine, liver, lung Treg.A displayed more transcriptional deviations. A conserved set of genes was upregulated, including *Nr4a1, Junb, Jund, Fos*, and *Dusp10*—genes associated with TCR signaling.

Thus, while Treg.A represents a broadly conserved resting Treg state present across lymphoid and non-lymphoid organs, its transcriptional profile is modulated in tissue-specific ways. These findings reveal the existence of intraparenchymal cluster A Tregs in NLTs and highlight some transcriptional adaptation across anatomical environments.

## A shared atlas of Treg states across tissues

A substantial body of work has described specialized Treg populations residing in NLTs^6,9,11,12,16,19,20,22,38–45^, prompting us to systematically examine the distribution of immgenT- defined Treg clusters across tissues under healthy conditions (**Fig. 4a**). All Treg clusters were detected across both lymphoid and non-lymphoid organs. Rather than being tissue-restricted, clusters differed primarily in their *relative abundance* across organs. Several reproducible distribution patterns emerged. Barrier tissues, including skin, small intestine, and colon, were enriched for Treg.D and E. Treg.E corresponded to RORγt⁺ Tregs (**Extended Data Fig. 5d,e**) and was most prominent in the colon, whereas Treg.D was enriched in the small intestine and skin (**Fig. 4b**). In contrast, Treg.F (discussed in detail below) showed a distinct pattern, being markedly enriched in tissues less commonly emphasized in Treg studies—such as kidney, liver, placenta, and mammary gland—where it comprised 22–77% of all Tregs.

**Figure 4.**
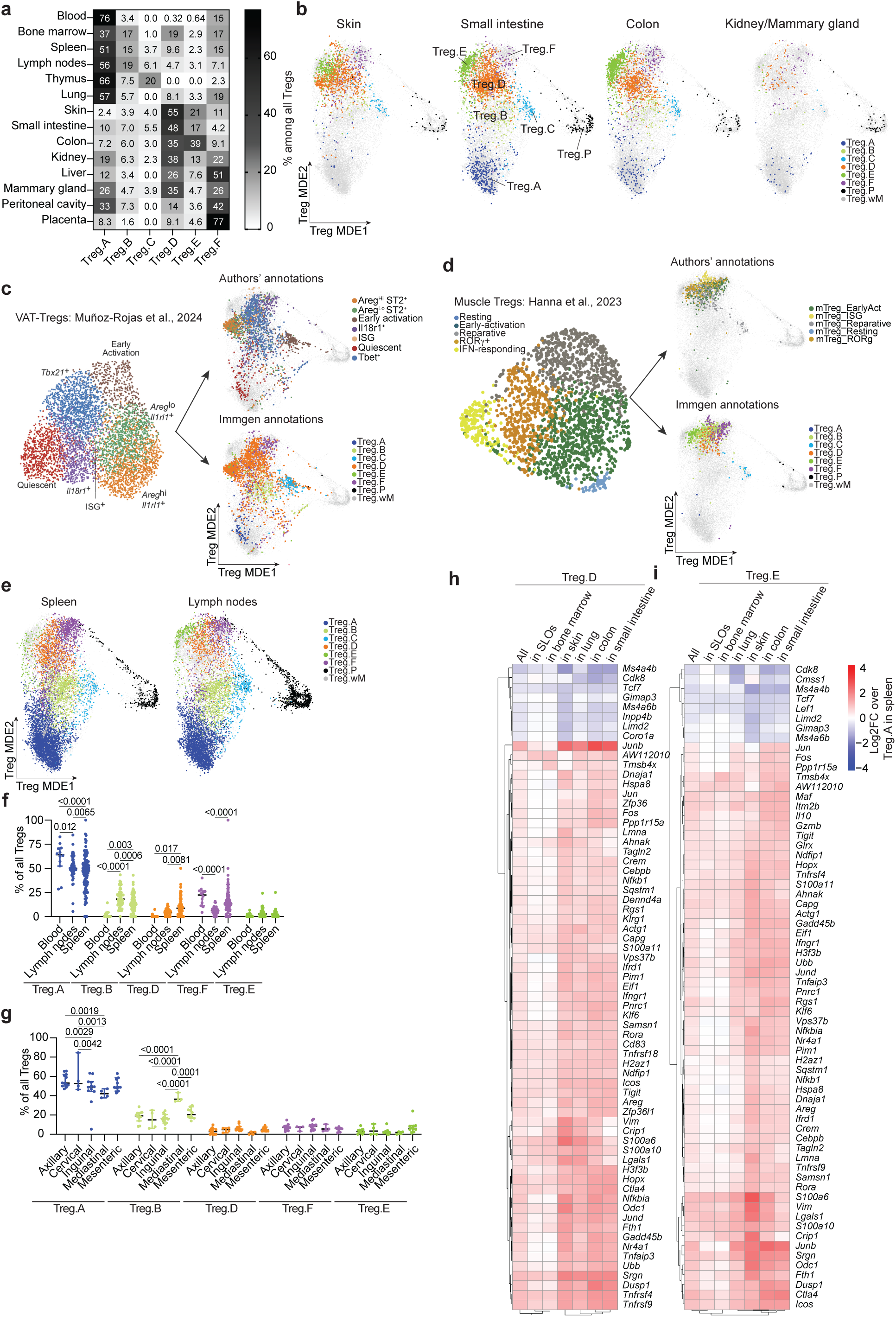
Treg cells in healthy tissues: shared clusters with organ-specific enrichments. **(a)** Heatmap showing the distribution of Treg clusters across healthy tissues in immgenT. Values represent median proportions among all Tregs (n = 1–60 samples per tissue). (b) Treg-MDE showing cluster distributions in healthy non-lymphoid tissues, including skin, small intestine, colon, and kidney and mammary gland. **(c,d)** T-RBI (T Reference-Based Integration) of published datasets from Muñoz-Rojas et al.^20^ **(c)** and Hanna et al.^9^ **(d).** Left, original authors’ annotations and UMAP embeddings. Right, projection of the same cells onto the immgenT Treg-MDE, shown with the original authors’ annotations (top) and immgenT cluster annotations (bottom). **(e)** Treg-MDE showing cluster distributions in healthy spleen and lymph nodes. **(f)** Treg cluster proportions in blood, lymph nodes, and spleen among all Tregs in immgenT samples. Summary statistics show median ± interquartile range. Statistical significance was assessed by two-way ANOVA with Tukey’s multiple-comparisons test. **(g)** Treg cluster proportions in axillary, cervical, inguinal, mediastinal, and mesenteric lymph nodes among all Tregs in immgenT samples. Summary statistics show median ± interquartile range. Statistical significance was assessed by two-way ANOVA with Tukey’s multiple-comparisons test. **(h)** Heatmap showing a shared transcriptional signature of Treg.D across organs. Values are shown as log2 fold change relative to Treg.A in the spleen. Genes shown meet fold-change criteria (>2 or <0.5) with adjusted p value < 0.05 in at least one tissue. **(i)** Heatmap showing a shared transcriptional signature of Treg.E across organs. Values are shown as log2 fold change relative to Treg.A in the spleen. Genes shown meet fold-change criteria (>2 or <0.5) with adjusted p value < 0.05 in at least one tissue. *ISG, interferon-stimulated genes; SLOs, secondary lymphoid organs*.

Because immgenT does not include certain well-characterized tissue Treg populations— such as those from visceral adipose tissue (VAT), skeletal muscle, and meninges—we next asked whether Tregs from these sites exhibit similar states. We applied reference-based integration (T- RBI), a custom algorithm introduced in detail in the accompanying immgenT-Cosmology manuscript^29^, which maps external single-cell datasets onto the immgenT reference. All three published datasets of tissue Tregs^9,16,20^ mapped robustly onto the immgenT Treg reference without forming new clusters (**Fig. 4c,d; Extended Data Fig. 6a,b**), indicating that the immgenT framework accounted for all major cell states across tissues. In VAT Tregs^20^ (**Fig. 4c**), immgenT annotations closely matched the original classifications: Areg⁺ST2⁺ cells mapped to Treg.D, quiescent VAT-Tregs to Treg.A, early activation states to Treg.D, IL18R1⁺ cells to Treg.F, and T-bet⁺ Tregs were distributed between Treg.B and a subset of Treg.D. Similarly, in skeletal muscle Tregs^9^ (**Fig. 4d**), RORγt⁺ cells mapped to Treg.F, resting Tregs to A, reparative Treg to D, and interferon-stimulated states to a subset of Treg.F. Additional rare populations, including Treg.B, Treg.F, proliferating Tregs (Treg.P), and Treg.C, were also detected, demonstrating that the immgenT framework captures the full diversity of VAT and muscle Treg states.

Effector Treg clusters typically associated with NLTs were also detectable in lymphoid tissues, albeit at lower frequencies (**Fig. 4e,f**). Notable differences were observed across blood, spleen, and lymph nodes, as well as among lymph nodes draining distinct anatomical sites. Treg.F (CD49b⁺) represented the dominant effector-like population in blood. Lymph nodes were themselves heterogeneous (**Fig. 4g; Extended Data Fig. 7a-e**): mediastinal lymph nodes were enriched for Treg.B, whereas mesenteric lymph nodes showed a relative increase in Treg.E, consistent with their drainage of gut-associated tissues.

To determine how conserved Treg states are transcriptionally across organs, we compared gene expression profiles of the same cluster across tissues. Using splenic Treg.A as a reference, we examined genes with FC > 2 or < 0.5 for Treg.E (**Fig. 4h**) and D (**Fig. 4i**) across organs. Core Treg.D and E signatures—including *Il10, Maf, Icos, Ctla4*, and *Rora*—were conserved across all tissues, including secondary lymphoid organs. In NLTs, these signatures were generally stronger, with additional upregulation of TCR signaling–associated genes for Treg.E and tissue-specific modulation for Treg.D (e.g., increased *Vim* and *Crip1* expression in skin cluster D and E).

In summary, healthy tissues harbor a shared repertoire of conserved Treg clusters, with relative abundance varying organ-specifically. While transcriptional identities are largely preserved across tissues, effector signatures are differentially amplified in NLTs. ImmgenT therefore provides a unified framework for dissecting both conserved and tissue-adapted features of Treg heterogeneity across the body.

### Treg during inflammation: redistribution of existing clusters rather than emergence of new states

Tregs perform multiple functions during inflammation, including maintaining tolerance to self-antigens, shaping effector and memory T cell differentiation, dampening inflammation, and promoting tissue repair^1,46^. We therefore asked how the Treg cluster distribution observed in healthy tissues (Fig. 4) changes across inflammatory settings, and whether inflammation drives the emergence of new Treg states.

Across all tissues and conditions, no inflammation-specific Treg clusters emerged. Instead, inflammation reshuffled the relative proportions of the main existing six clusters in tissue- and context-specific ways (**Fig. 5a; Extended Data Fig. 8a**). In the spleen, most conditions closely resembled baseline, with Treg.A remaining dominant, whereas LCMV clone 13, NP–OVA immunization, and *T. gondii* infection led to expansion of Treg.D–F effector clusters. In the colon, inflammation broadly increased Treg.F (and, to a lesser extent, Treg.B) at the expense of Rorc^+^ Treg.E, with *C. rodentium* additionally inducing a strong proliferative response. In the uterus, where Treg.F predominates at baseline, Chlamydia infection and SFB colonization shifted the balance toward Treg.D and E. The lung was also dynamic: at baseline it contained very few Treg.D, B, or E cells, but inflammatory exposures drove pronounced polarization. House dust mite (HDM)-induced allergy favored a Treg.E bias with clear *Rorc* expression (**Fig. 5b**), whereas influenza and tuberculosis increased Treg.D, marked by *Klrg1* expression (**Fig. 5c**), similar to the ones seen traditionally in healthy colon (**Extended Data Fig. 8b,c**).

**Figure 5.**
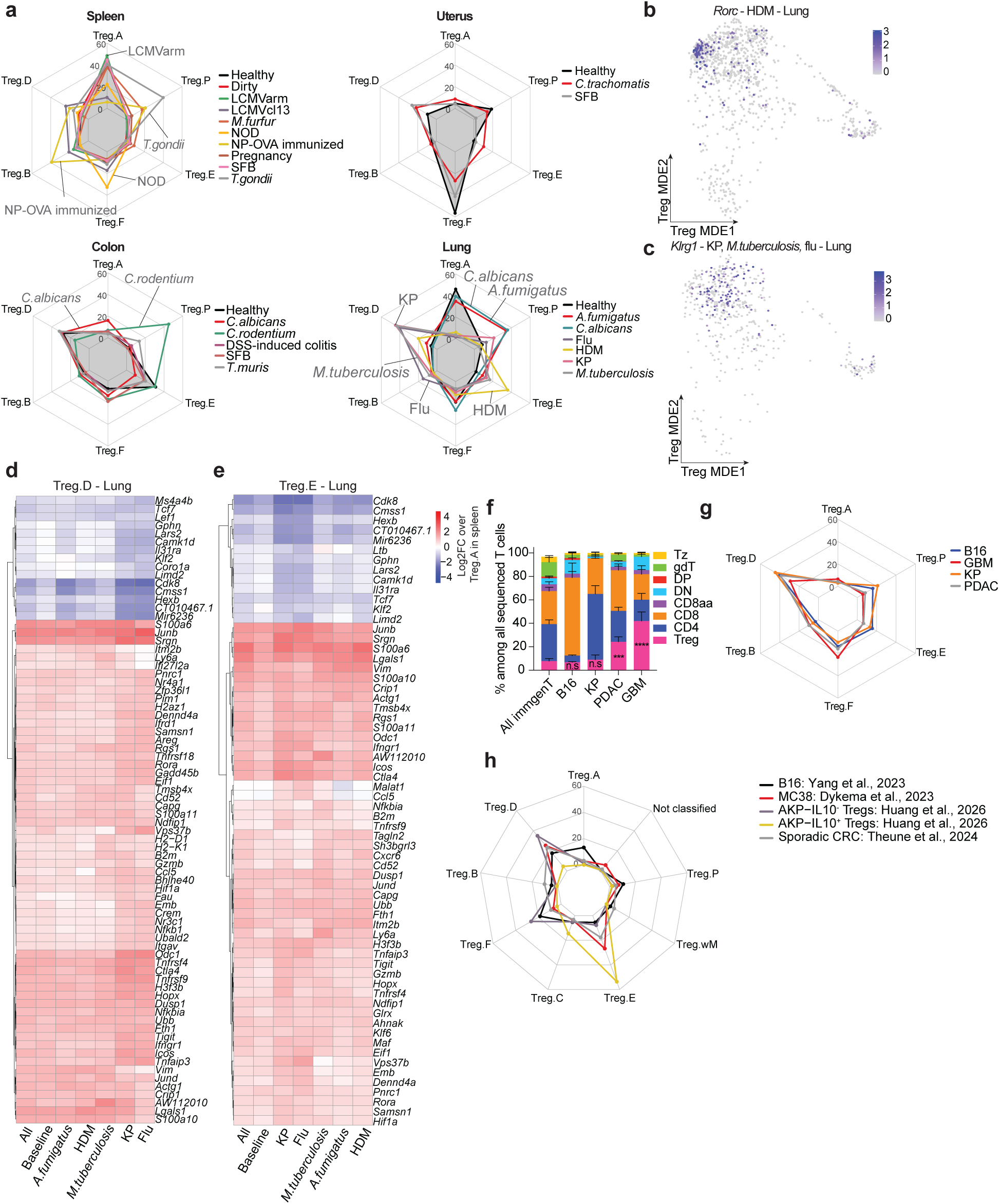
Treg cells during inflammation in tissues undergo cluster redistribution rather than the emergence of new states. **(a)** Radar plots showing Treg cluster distributions across immune perturbations compared to baseline (healthy, black) in spleen, uterus, colon, and lung. Values represent median proportions. (b) Treg-MDE showing expression of *Rorc* in lung Tregs following house dust mite (HDM) exposure. **(c)** Treg-MDE showing expression of *Il1rl1* in lung Tregs from KP lung cancer, *Mycobacterium tuberculosis*, and influenza infection (combined). **(d)** Heatmap showing a maintained transcriptional signature of Treg.D in the lung across immune perturbations. Values are shown as log2 fold change relative to Treg.A in the spleen. All genes with fold-change >2 or <0.5 and adjusted p value < 0.05 in at least one condition. **(e)** Heatmap showing a maintained transcriptional signature of Treg.E in the lung across immune perturbations. Values are shown as log2 fold change relative to Treg.A in the spleen. All genes with fold-change >2 or <0.5 and adjusted p value < 0.05 in at least one condition. **(f)** Stacked bar plots showing the proportion of Tregs among total T cells in different cancers (B16 melanoma, glioblastoma (GBM), KP lung cancer, and orthotopic PDAC) profiled in immgenT, compared to the average proportion across the full dataset. Values are shown as median ± SEM. Statistical significance by two-way ANOVA with Dunnett’s multiple-comparisons test versus all non-tumor tissues (n.s., p > 0.05; *** p = 0.0003; **** p < 0.0001). **(g)** Radar plots showing Treg cluster distributions in the same cancers as in (f). Values represent median proportions. **(h)** Radar plots showing Treg cluster distributions in external tumor datasets mapped with T-RBI: B16 melanoma^48^, MC38 (WT and MC38-GP) colorectal cancer^49^, IL-10^-^ and IL-10^+^ Tregs from orthotopic AKP tumor^50^, and sporadic CRC^51^. Values represent median proportions. *LCMVarm: LCMV armstrong; LCMVcl13: LCMV clone 13; NOD: Non-Obese Diabetic; NP- OVA: nitrophenyl-conjugated ovalbumin; SFB: Segmented Filamentous Bacteria; DSS: Dextran Sulfate Sodium; HDM: House Dust Mite; KP: activation of the KrasG12D oncogene and deletion of the p53 tumor suppressor; GBM: Glioblastoma; PDAC: Pancreatic Ductal Adenocarcinoma; AKP tumor: mutations in Apc, Kras, and p53 genes; MC38-GP: MC38 cancer cell line expressing the LCMV glycoprotein (GP)*.

We next asked whether inflammation alters transcriptional identity in addition to reshaping cluster proportions. Focusing on lung Treg.D and F (**Fig. 5d,e**), we compared each inflammatory condition to splenic Treg.A as a reference and identified genes with FC > 2 or < 0.5 and adjusted *P* < 0.05. Most differentially expressed genes across conditions belonged to the original cluster signatures, indicating preserved transcriptional identity. Differences largely reflected quantitative changes in expression levels rather than gain or loss of signature genes. For Treg.D, influenza enhanced expression of genes such as *Ifngr1, Tnfrsf9* (4-1BB), *Tnfrsf4* (OX40), *Ctla4, Icos,* and *Tigit*. For Treg.E, inflammation amplified components of the canonical signature, including *Ifngr1, Icos, Ctla4, Lgals1,* and *S100a6*. Thus, inflammation did not generate new Treg clusters. Instead, it altered the relative frequencies of existing clusters—sometimes substantially compared to baseline—and modulated expression levels within their established transcriptional programs.

We next investigated Treg cluster distribution in cancer. Tumor infiltrating Tregs have been widely studied in different cancer types^47^. Some tumors were highly enriched for Tregs— from 7% of all T cells in B16 to 42% in glioblastoma (GBM) (compared to 8% across all immgenT samples; **Fig. 5f**). Treg.D dominated tumor Tregs (30-45%) with some enrichment in Treg.F and E (**Fig. 5g**) but no apparent systematic changes upon cancer treatments (**Extended Data Fig. 9a**). Tumor models are widely heterogeneous so we used T-RBI to extend our analysis to external datasets covering B16 melanoma^48^ and other cancers such MC38 colorectal cancer^49^, orthotopic AKP (colorectal cancer model characterized by mutations of Apc, Kras, and P53 genes) tumor^50^, and sporadic colorectal cancer^51^ (**Extended Data Fig. 9b-e**). In total, we added 28,909 Treg cells to our analysis. While Treg.D was well represented across all tumor models (22% in B16, 32% in MC38, 44% in AKP IL-10^+^ Tregs, and 6% in AKP IL10^-^ Tregs), Treg.E was selectively overrepresented in the MC38 model (33%) and AKP tumor IL-10^+^ Tregs (67%; **Fig 5h, Extended Data Fig. 9f**). These findings confirm a general preference for Treg.D across cancers, where most *Ccr8^+^* Tregs map (**Extended Data Fig. 9g)**, with differential representation of other Treg clusters depending on tumor origin. Of note, less than 8% of cells are not annotated but mapped to the Treg space and had a low discovery score confirming that our framework captured the spectrum of Treg states in cancer (**Fig. 5h, Extended Data Fig. 9f**).

### Treg.F: a circulating effector Treg population enriched in specific NLTs and characterized by CD49b and *Klf2* expression

Among the eight major Treg clusters identified in the immgenT atlas, Treg.F emerged as an unexpected and distinct population. First, Treg.F was markedly enriched in specific tissues: placenta (65%), peritoneal cavity (42%), liver (31%), and uterus (56%) and blood (15%) (**Fig. 6a,b**). In the placenta, Treg.F was particularly prominent, comprising 60% of all Tregs (corresponding to 0.8% of total T cells), and it was also detectable in the uterus of virgin mice, albeit at lower overall frequency. In the circulation, Treg.F represented the only CD44⁺ Treg population and thus constituted the dominant circulating effector Treg compartment (**Fig. 4a,e**). To exclude the possibility that Treg.F detection was due to blood contamination, we performed intravascular labeling (**Extended Data Fig. 10a**). Extravascular CD44⁺CD49b⁺ Tregs in both the spleen and liver were present, supporting *bona fide* parenchymal Treg.F cells.

Second, the transcriptional program of Treg.F uniquely combined activation-associated genes (*S100a10, S100a11*) with genes involved in migration and tissue trafficking (**Fig. 6c,d; Extended Data Fig. 10b**), including *Klf2*, *Itgb1, Itgb7,* and *Itga2*. CITE-seq analysis revealed that this population was marked at the protein level by CD29 (*Itgb1*) and CD49b (*Itga2*), despite relatively modest differential expression of *Itga2* transcripts, underscoring the imperfect correspondence between integrin mRNA and surface protein abundance. These features closely parallel the CD49b⁺ Tregs previously described by Fan et al.^52^ Notably, CD29 alone was insufficient to define this population, consistent with its ability to pair with multiple integrin α- chains; gating on CD29⁺CD49b⁻ cells failed to recover Treg.F in CITE-seq (**Extended Data Fig. 10c**), indicating that CD49b is required for specific identification by flow cytometry.

Third, one of the most strongly enriched genes in Treg.F was the transcription factor *Klf2* (**Fig. 6f,g**). Although *Klf2* is expressed in resting Tregs (e.g., Treg.A), it was selectively upregulated in Treg.F compared with other effector Treg populations. *Klf2* plays a central role in T cell migration^53–55^, in part through regulation of *S1pr1*, enabling recirculation through the blood (**Fig. 6h,i**). Consistent with this model, *S1pr1* expression among effector Tregs was largely restricted to Treg.F, and prior work showed that FTY720-mediated blockade of S1PR1 abrogates circulating CD49b⁺ Tregs^52^. *Klf2*-specific deletion in Tregs, however, did not decrease CD49b⁺CD29⁺ Tregs in the liver and was associated with an increased frequency of these cells in the spleen (**Extended Data Fig. 10d,e**), indicating that *Klf2* is not required for the generation or maintenance of Treg.F and that its effects on this cluster may be tissue dependent.

Finally, Treg.F was not confined to homeostasis or pregnancy. This population was also present and often enriched in inflammatory contexts, including Chlamydia infection in the uterus, cancer models following immune checkpoint inhibition, and in NOD mice (**Fig. 6j**).

In summary, Treg.F represents the principal circulating effector-like Treg population and shows preferential enrichment in a subset of non-lymphoid tissues. Defined by CD49b expression and selective upregulation of *Klf2*, this state seems to integrate activation and migratory programs consistent with S1PR1-dependent recirculation. Together, these features position Treg.F as a distinct and functionally meaningful Treg state.

### Treg cells in MHC-II-deficient mice

Splenic CD4^+^ T cells from Aβ^o/o^ mice deficient in MHC-II molecules were profiled in order to identify unconventional CD4 T cells restricted by other MHC-like molecules (IGT90/91). While the heterozygous littermates showed the dominant CD4^+^ lineage, along with some Treg and a few PLZF⁺ Tz cells MHC-II-deficient homozygotes showed a massive drop of conventional CD4^+^ T cells, with a compensating expansion of Tz cells (**Fig. 7a-c**), as expected^56,57^. Interestingly, a sizeable population of FoxP3⁺ cells also persisted at high frequency (**Fig. 7a-b**), in hindsight not unexpectedly given prior reports of Treg-like cells in MHC-II deficient mice^58,59^. These Treg cells expressed FoxP3 (**Fig. 7c**) and mapped to the location of normal Treg cells on the MDE (**Fig. 7a**). Their distribution across Treg clusters was also comparable to normal Tregs, albeit with a surfeit of Treg.F (**Extended Data Fig. 11a**). αβTCR sequencing confirmed that Treg cells from MHC-II deficient mice did not express CD1-restricted TCRs typical of iNKT or MAIT cells. They did display more clonal expansions relative to proficient littermates (**Fig 7d**), and TRAV and TRBV usage generally aligned with those of normal Treg cells (**Extended Data Fig. 11b**). Flow cytometry confirmed the presence of CD4^+^FoxP3^+^ cells in Aβ^o/o^ mice (**Fig. 7e**), with reduced CD25 expression. This was confirmed in mice with deletion of both MHC-II beta chain genes^60^, ruling out selection by an AαEβ that is theoretically possible in Aβ^o/o^ mice (**Fig. 7e**). Absolute Treg numbers were reduced by approximately an order of magnitude in spleen, lymph nodes, and gut (**Fig. 7f**). In the colon lamina propria, the dominant population of RORγ^+^ Tregs was completely lost in the absence of MHC-II (**Fig. 7g)**, leaving only the Helios^+^ compartment.

**Figure 6.**
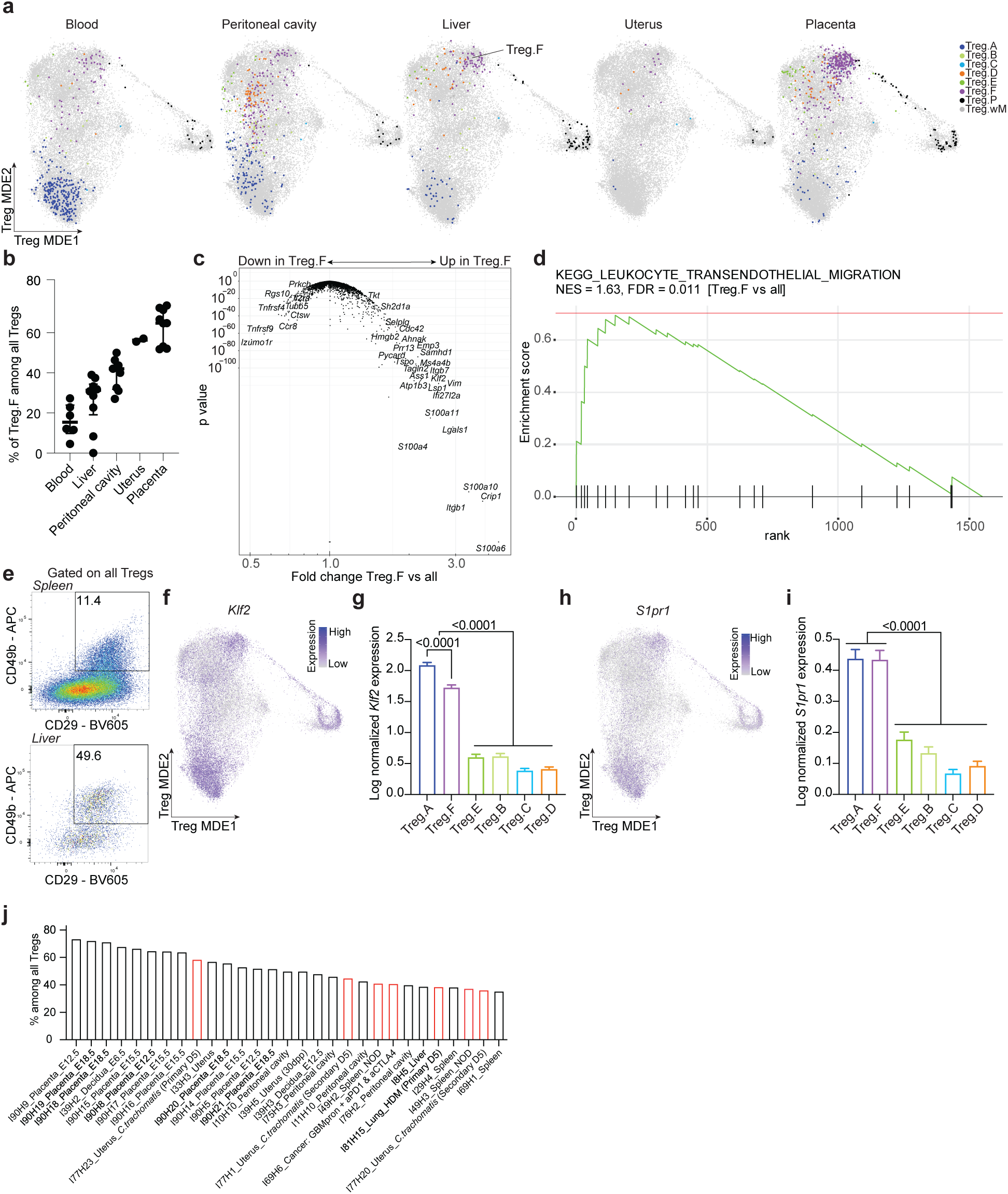
Treg.F: the main circulating effector Treg population, enriched in specific non- lymphoid tissues and characterized by CD49b and KLF2 expression. (a) Treg-MDE showing Treg cluster distributions in blood, peritoneal cavity, liver, uterus, and placenta from healthy samples. (b) Proportion of Treg.F among all Tregs in blood, peritoneal cavity, liver, uterus, and placenta. Summary statistics are shown as median values with interquartile range. (c) Volcano plot showing differential gene expression between Treg.F and all other Treg clusters, excluding proliferating and miniverse clusters. Values are shown as fold change versus *P* value. (d) Gene set enrichment analysis (GSEA) of the Treg.F transcriptional signature derived in (c). The most significantly enriched pathway is shown (KEGG leukocyte transendothelial migration; NES = 1.63, FDR = 0.011). (e) Representative flow cytometry plots showing CD29 and CD49b expression on Tregs in spleen and liver. Data are representative of 3–4 independent experiments. (f) Treg-MDE showing expression of *Klf2*. (g) *Klf2* expression (log1p CP10K) across Treg clusters. Values are shown as median ± SEM. Statistical significance was assessed by one-way ANOVA with Tukey’s multiple-comparisons test. (h) Treg-MDE showing expression of *S1pr1*. (i) *S1pr1* expression (log1p CP10K) across Treg clusters. Values are shown as median ± SEM. Statistical significance was assessed by one-way ANOVA with Tukey’s multiple-comparisons test. (j) Ranked bar plot showing the top 25 samples with the highest frequency of Treg.F among all Tregs. Healthy samples are shown in black and immune perturbations in red. Sample identifiers correspond to immgenT sample IDs. *NOD: Non-Obese Diabetic; HDM: House Dust Mite; GBM: Glioblastoma*.

**Figure 7.**
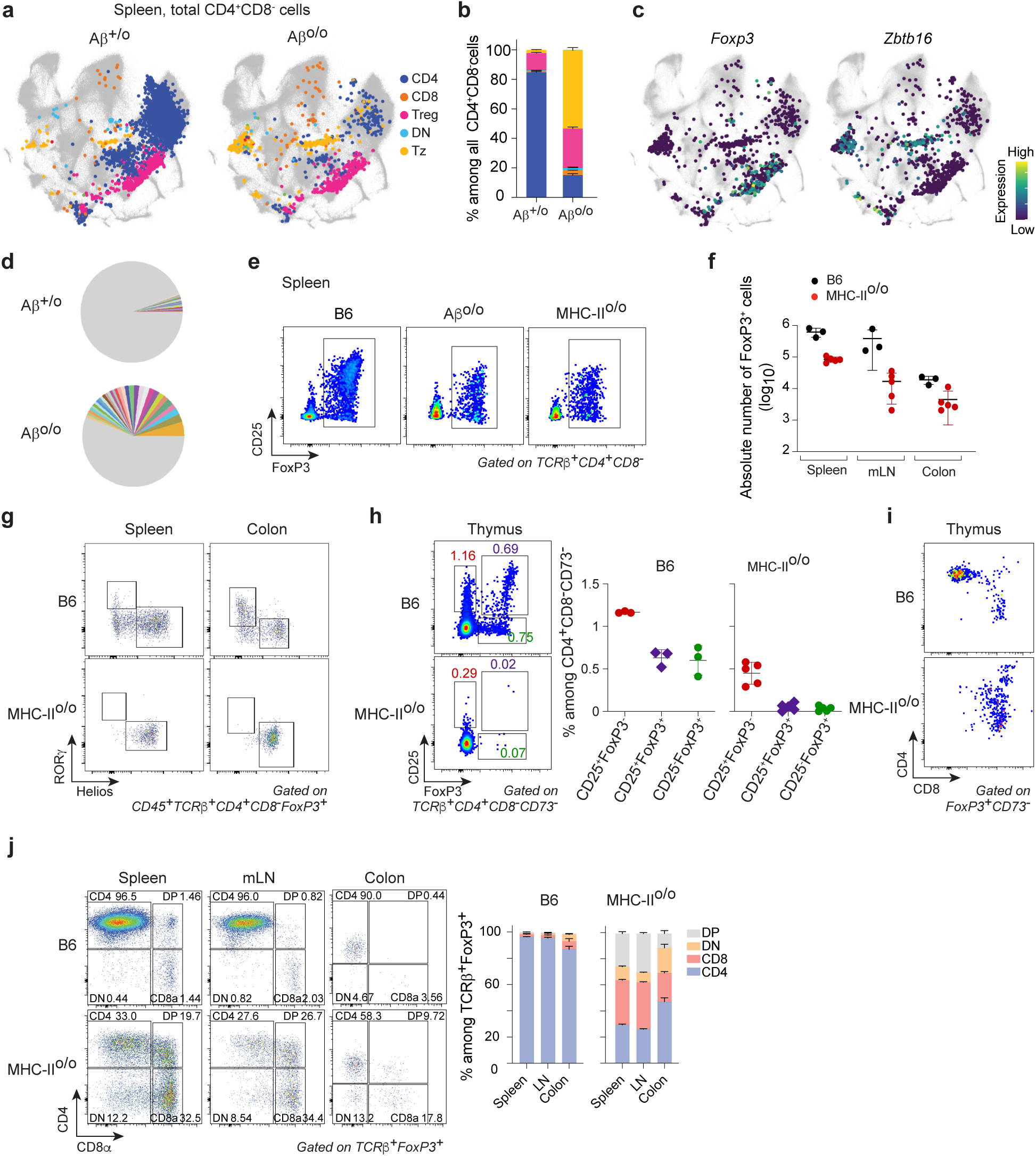
Persistence of Treg cells in MHC-II-deficient mice. (a,b) Lineage distribution of total CD4^+^ CD8^-^ cells in Aβ^+/o^ and Aβ^o/o^ mice. **(a)** Treg-MDE plot with cells colored by lineage. **(b)** Quantification shown as a stacked bar plot. Aβ^+/o^, n=6; Aβ^o/o^ n=2, IGT91/92). (c) Treg-MDE showing the expression of *Foxp3* and *Zbtb16* in the same Aβ^o/o^ samples as in (a,b). **(d)** Pie charts showing the frequencies of TCR clonotypes in Aβ^+/o^ and Aβ^o/o^ mice (same samples as in a,b). Gray indicates clonotypes represented by a single cell. **(e)** Representative flow cytometry plots of CD25 and FoxP3 expression gated on spleen TCRβ^+^CD4^+^CD8^-^ from B6, Aβ^o/o^ and MHC-II^o/o^ mice. **(f)** Absolute numbers (log10) of FoxP3^+^ cells in the spleen, mesenteric lymph nodes (mLNs), and colon of B6 and MHC-II^o/o^ mice. Results are shown as medians with 95% confidence intervals from 4 independent experiments. **(g)** Representative flow cytometry plots of RORγ and Helios expression gated on CD45^+^TCRβ^+^CD4^+^CD8α^-^FoxP3^+^ cells from the spleen and colon of B6 and MHC-II^o/o^ mice. **(h)** (Left) Representative flow cytometry plots of CD25 and FoxP3 expression gated on thymic TCRβ^+^CD4^+^CD8^-^CD73^-^ cells from B6 and MHC-II^o/o^ mice. (Right) Proportions of CD25^+^FoxP3^-^, CD25^+^FoxP3^+^, and CD25^-^FoxP3^+^ cells among CD4^+^CD8^-^CD73^-^ cells in the thymus of B6 and MHC-II^o/o^ mice. **(i)** Representative flow cytometry plots of CD4 and CD8 expression gated on thymic FoxP3^+^CD73^-^ cells from B6 and MHC-II^o/o^ mice. **(j)** (Left) Representative flow cytometry plots of CD4 and CD8α expression in TCRβ^+^FoxP3^+^ cells from the spleen, mLNs, and colon of B6 and MHC-II^o/o^ mice. (Right) Stacked bar plot showing the proportions of DP (CD4^+^CD8α^+^), DN (CD4^-^CD8α^-^), CD4^+^CD8α^-^, and CD4^-^CD8α^+^ among TCRβ^+^FoxP3^+^ cells in the indicated tissues from B6 and MHC-II^o/o^ mice (n = 3–6). Results are shown as median with interquartile range from 3 independent experiments.

Prior studies have reported the presence of FoxP3⁺ cells in MHC-II-deficient settings and proposed that they arise from peripheral differentiation (pTreg) rather than thymic selection^58^. To address this question, we examined thymic Treg cells and their CD25^+^ or FoxP3^+^ single positive precursors^61,62^. In the absence of MHC-II, mature CD25^+^FoxP3^+^ thymocytes were very severely reduced, as were the FoxP3^+^ single-positive precursors, while the CD25^+^ precursor population was less affected (**Fig. 7h**). Gating on FoxP3^+^ TCRβ^+^ cells showed that ∼90% of TCRβ⁺FoxP3⁺ cells were CD4⁺ single-positive (CD4SP), with ∼5% CD8⁺ SP in WT mice (**Fig. 7i**). In MHCII^o/o^ thymi, the mature CD4SP population of FoxP3^+^ cells was essentially absent, leaving only CD4^dull^CD8^dull^intermediates and CD8SP. Thus, in the absence of MHC-II molecules, differentiation of FoxP3^+^ cells towards CD4^+^ Tregs seems to abort, leaving only the CD8SP branch.

In the periphery, the CD4/CD8 distribution of FoxP3^+^ cells partially reflected the thymus, with marked presence of CD4^+^CD8^+^ and CD8^+^ cells, considerably increased in proportion relative to the heterozygous control (**Fig. 7j**). The latter likely correspond to the small group of FoxP3^+^ CD8^+^ Treg cells that have been reported recently, and whose regulatory or cytotoxic function remains debated^63,64^. Note that FoxP3^+^CD8^+^ cells did not appear in sizable numbers in any of the challenged immgenT datasets, and seem to be most revealed by the MHC-II deficiency, perhaps reflecting a competition by CD4^+^ Tregs for differentiation in the thymus (**Fig. 7h**). As to the CD4^+^ FoxP3^+^ component, consistent with **Fig. 7a,e**, we can’t distinguish whether it stems for peripheral Treg differentiation (pTreg) or from homeostatically-driven expansion of the few Treg cells that differentiate in the thymus (consistent with their restricted repertoire; **Fig. 7d**).

In summary MHC-II deficient mice retain several populations of FoxP3⁺ suppressive cells. Their restriction elements are unknown at this point, but they transcriptionally overlap with WT Tregs and their presence likely explains why MHC-II deficient mice and patients do not develop systemic autoimmunity.

## Discussion

A large and diverse array of Treg subsets has been described over the past two decades, defined by developmental origin, surface phenotype, tissue localization, and functional specialization^20,33,65^. However, how these populations relate to one another at the molecular level has remained unclear. ImmgenT provides a unifying framework that places previously described Treg populations into a coherent molecular landscape (**Extended Data Table 4**).

The strength of our analysis lies in the breadth and diversity of conditions profiled, encompassing lymphoid organs, multiple non-lymphoid tissues at baseline, and a wide range of immune perturbations. To assess the completeness of this framework, we applied reference-based integration (T-RBI) to multiple external single-cell datasets profiling Tregs from contexts not represented in immgenT, including visceral adipose tissue^20^, skeletal muscle^9^, meninges^16^, and tumors^48–51,66^. In all cases, Tregs mapped robustly onto the existing immgenT state space without forming additional clusters, supporting the conclusion that immgenT-Treg captures the major axes of Treg heterogeneity in the mouse.

Treg diversity resolves into a relatively simple organization structured around eight conserved transcriptional clusters. These clusters correspond to familiar and previously described Treg states, yet the convergence of a diverse array of reported Treg subsets into a limited number of robust clusters—and the particular states that emerge as dominant organizers of the landscape— was to some extent surprising. The dominant axis separates resting from activated Tregs, reflecting long-standing distinctions^17,19^. Effector states include clusters corresponding to well-known KLRG1⁺ ST2⁺ Tregs (Treg.D)^20,38,43,67–69^, RORγt⁺ Tregs (Treg.E)^11,33,44,70^, and Treg.B (which includes T follicular regulatory cells)^25,71–73^. Treg.C represents a frequently observed but less well-characterized state marked by early TCR signaling^9,17,20^ and apparently limited effector specialization, reminiscent of the “undifferentiated” activated CD44⁺ CD4 T cells (as described in the accompanying immgenT-CD4 manuscript^74^).

One of the most unexpected findings was the prominence of Treg.F, which unifies previously described CD49b⁺ ^52^ and Klf2⁺ ^53,54^ effector Tregs into a single, distinct cell state. Although Treg.F is enriched in highly vascular tissues such as placenta and liver, vascularity alone does not explain its distribution^52^, as it is absent from the kidney and not particularly enriched in spleen or lung despite their dense vasculature. Notably, while the majority of circulating Tregs are resting, the activated fraction in blood is nearly exclusively composed of Treg.F, consistent with its high expression of *Klf2* and *S1pr1*. These findings may have implications for human studies that rely heavily on blood sampling. The functional role of Treg.F remains to be fully elucidated, particularly in the placenta, where it is most prominent and may contribute to fetal tolerance or other pregnancy-associated immune adaptations.

Across organs, Treg heterogeneity was largely preserved, with all major clusters present but differentially enriched depending on the organ, consistent with previous reports^9,20,75^ . Barrier tissues preferentially contained Treg.E (RORγt⁺) and Treg.D (KLRG1⁺ ST2⁺) effector states, whereas lymphoid organs were enriched in resting Tregs (Treg.A). Our analysis highlights tissues less studied in Treg biology, such as the liver, peritoneal cavity, and placenta, which were all enriched for Treg.F, as discussed above. In parallel, the presence of resting-like Treg cells in non-lymphoid tissues raises the possibility that some may reside in tertiary lymphoid structures and serve as a local reservoir.

Inflammatory settings have often been used to define additional Treg subsets, such as T-bet⁺ Tregs^18,23,24,65^. Contrary to expectations, no new cluster emerged under inflammatory conditions that were not already present at baseline. Instead, inflammation altered the relative abundance of existing clusters, potentially through mechanisms including plasticity, differential recruitment, proliferation or survival. T-bet expression did not define a discrete cluster in the immgenT data; instead, it appeared expressed across multiple effector clusters. This observation is consistent with our analysis of Th1 cells in the immgenT-CD4 companion manuscript^74^, in which Th1 cells were distributed across four clusters with varying degrees of polarization, including overlaps with other Th programs. More broadly, T-bet was also widely expressed across activated cells from all T cell lineages, as described in the immgenT-Cosmo manuscript^29^. It does not negate the functional importance of T-bet in Tregs, as genetic ablation studies demonstrate non-redundant roles^18^, but suggests that T-bet⁺ Tregs are heterogeneous. More broadly, inflammation primarily altered the relative abundance of conserved Treg states. These findings raise questions about the context-specific roles of individual clusters, such as Treg.E in house dust mite–induced allergic inflammation (HDM). They also prompt investigation into how different immune challenges shift Treg transcriptional states within a given tissue (e.g., through plasticity or migration), and why multiple activated Treg states coexist within the same tissue—whether they fulfill distinct or complementary functions and occupy different niches.

Importantly, this organization does not imply an absence of variation within clusters. For example, Treg.D cells from different conditions do not map to exactly the same location in the state space, highlighting additional layers of transcriptional heterogeneity. Complementary gene program (GP) analyses within immgenT (immgenT-GP manuscript) further capture this variation, supporting a model in which Tregs occupy a continuum of gene program expression^17,19,76^.

In conclusion, because scRNAseq datasets and routine flow cytometry data can relate directly to the immgenT framework, immgenT-Treg can serve as a common reference to define and compare Treg identity across studies (see Methods - immgenT resource: public data browsers). Rather than a static classification, it is designed as a practical and reusable resource. The immgenT-Treg flow cytometry panel, partly overlapping with previously published strategies^13,42,52,77,78^, bridges conventional flow cytometry and high-dimensional single-cell transcriptomics, providing a uniform approach to project flow-based data onto the immgenT space. Additional flow cytometry markers tailored to specific biological questions, such as T-bet staining to identify cells expressing a Th1-like program, could further complement this panel. In parallel, T-RBI enables mapping of external single-cell datasets onto the atlas, facilitating direct comparisons across studies. Together, immgenT-Treg establishes a common framework for Treg heterogeneity and equips the field with a foundation for future discovery.

## Supporting information

Extended Data Table 1 - Treg Samples and Metadata

Extended Data Table 2 - Treg Cluster Signatures

Extended Data Table 3 - immgenT Treg flow cytometry panel

Extended Data Table 4 - immgenT Previous Nomenclature

## Acknowledgments

We thank the many colleagues who were consulted at various stages of this project. We thank Dr. Stephen C. Jameson (University of Minnesota) for sharing the Klf2^fl/fl^ mouse strain used in this study. This work was funded by a grant from the NIH to the ImmGen consortium (R24-072073).

## Author contributions

AF, XC, JAM, IM performed the experiments; AF, DZ, NM, IA, CO, IM, XC, SZ, and CJI participated in the analysis; AF, DZ, JRH, LB, DV and CB designed the overall study; all principal investigators provided funding and oversaw the experiments; AF and DZ wrote the manuscript with input from other authors.

## Collaborators

Participants in the immgenT Project include:

Aaron Liu^1^, Alexander Chervonsky^2^, Alexandra Cassano^2^, Alia Welsh^3^, Amir Ferry^11^, Ananda Goldrath^11^, Andrea Lebron-Figueroa^5^, Ankit Malik^2^, Anna-Maria Globig^4^, Antoine Freuchet^2^, Bana Jabri^2^, Charlotte Imianowski^6^, Christophe Benoist^5^, Claire Thefaine^7^, Dan Kaplan^6^, Dania Mallah^5^, Dario Vignali^6^, David Sinclair^5^, David Zemmour^2^, Derek Bangs^8^, Domenic Abbondanza^2^, Enxhi Ferraj^9^, Eric Weiss^6^, Erin Lucas^7^, Evelyn Chang^9^, Gavyn Chern Wei Bee^10^, Giovanni Galletti^11^, Ian Magill^5^, Iliyan D Iliev^12^, Joonsoo Kang^9^, Jordan Voisine^2^, Josh Choi^5^, Julia Merkenschlager^13^, Jun R. Huh^5^, Katharine Block^7^, Ken Cadwell^10^, Kennidy K. Takehara^11^, Kevin Osum^7^, Laurent Brossay^14^, Laurent Gapin^15^, Liang Yang^5^, Lizzie Garcia-Rivera^1^, Marc K. Jenkins^7^, Maria Brbic^16^, Maria-Luisa Alegre^2^, Marion Pepper^8^, Mariya London^17^, Matthew Stephens^2^, Maurizio Fiusco^16^, Melanie Vacchio^3^, Michael Starnbach^5^, Michel Nussenzweig^13^, Mitch Kronenberg^18^, Myriam Croze^19^, Nalat Siwapornchai^5^, Nathan Morris^12^, Nicole E. Scharping^11^, Nika Abdollahi^19^, Nitya Mehrotra^2^, Odhran Casey^5^, Olga Barreiro del Rio^5^, Paul Thomas^20^, Peter Carbonetto^2^, Remy Bosselut^3^, Rocky Lai^9^, Sam Behar^9^, Sam Borys^14^, Sara E. Hamilton^7^, Sara Mostafavi^8^, Sara Quon^11^, Serge Candéias^21^, Shanelle Reilly^14^, Shanshan Zhang^5^, Siba Smarak Panigrahi^16^, Sofia Kossida^19^, Stefan Muljo^3^, Stefan Schattgen^20^, Stefani Spranger^22^, Steve Jameson^7^, Susan M. Kaech^1^, Takato Kusakabe^12^, Taylor Heim^22^, Tianze Wang^8^, Tomoyo Shinkawa^9^, Ulrich von Andrian^5^, Val Piekarsa^5^, Véronique Giudicelli^19^, Vijay Kuchroo^5^, Woan-Yu Lin^12^, Ziang Zhang^2^

1. NOMIS Center, Salk Institute for Biological Sciences, 2. The University of Chicago, 3. National Institutes of Health, 4. Allen Institute for Immunology, 5. Harvard Medical School, 6. Dept of Dermatology and Immunology, University of Pittsburgh, 7. University of Minnesota, 8. University of Washington, 9. UMass Chan Medical School, 10. University of Pennsylvania, 11. University of California San Diego, 12. Weill Cornell Medicine, 13. The Rockefeller University, 14. Brown University, 15. University of Colorado Anschutz Medical Campus, 16. Swiss Federal Institute of Technology, Lausanne, 17. New York University, 18. La Jolla Institute, 19. IMGT, Univ Montpellier, 20. St. Jude Children’s Research Hospital, 21. Alternative Energies and Atomic Energy Commission, Grenoble, 22. Massachusetts Institute of Technology

## Methods

### Animals

Mice used in the immgenT dataset are described in detail in the immgenT Cosmology manuscript^29^ (Extended Data Table 1) and on the immgenT website (https://immgen.org/ImmGenT/), including sex, age, and genetic background. With rare exceptions, experiments were performed using C57BL/6 (B6) mice, most of which were sourced from The Jackson Laboratory. Experimental conditions, including infection models, immunization strategies, and tissue processing protocols, are detailed in the immgenT Cosmology manuscript^29^ (Extended Data Table 1), the GEO GSE297097 dataset, and the immgenT website. Both male and female mice were used.

The flow cytometry experiments presented in the present study were conducted on mice: Foxp3^EGFP^: JAX:006769, Foxp3-Ires-Thy1.1/B6^78^, Aβ° (H2-Aβ1tm1.Doi)^79^, MHC-II^-^: JAX:003584), and Foxp3^cre^Klf2^fl/fl^ mice. Klf2^fl/fl^ were originally generated^80^ by Dr. Stephen C. Jameson (University of Minnesota) and crossed to Foxp3-Cre mice by Dr. Laurent Brossay (Brown University), who kindly provided the resulting Foxp3^cre^Klf2^f/fl^. All mice were bred and maintained under specific pathogen–free conditions at the University of Chicago, in accordance with procedures reviewed and approved by the University of Chicago Institutional Animal Care and Use Committee (IACUC) guidelines (Protocol ID: 72714).

### immgenT dataset: experiments and data processing

The immgenT dataset comprises 66 experiments of single-cell RNA-seq, CITE-seq (128- plex), and paired TCRαβ sequencing (10x Genomics 5′ v2 platform), corresponding to 80 encapsulation runs. Each encapsulation is assigned a unique “IGTx” identifier (IGT1–96) used to track datasets. In a minority of cases, two IGT identifiers denote parallel encapsulations of the same biological samples. Individual cells are indexed by a unique IGT.cellID, and samples are tracked using hashtag identifiers (IGT.HT).

A detailed description of data acquisition and processing is provided in the immgenT Cosmology manuscript^29^. Briefly:

#### Logistics

Experiments were conducted across multiple laboratories in the United States. Participating laboratories carried out mouse treatments (e.g., infection or immunization) and tissue processing with their established materials and reagents. The immgenT research assistant (IM) traveled to the site, helped with sample CITE-seq labeling and cell sorting, and performed encapsulation and library preparation. Each experiment typically included ∼10 hashtagged and pooled samples, and a standardized spleen control for batch-effect assessment. Library construction and sequencing were centralized at the Broad Institute.

#### Sample processing, library construction, and sequencing

Flow cytometry–sorted T cells were multiplexed using TotalSeq-C hashtag^81^ antibodies (BioLegend), enabling pooling prior to encapsulation. Cells were stained with a custom 128-antibody TotalSeq-C panel and subjected to joint single-cell RNA, surface protein (CITE-seq), and TCR sequencing using the 10x Genomics 5′ v2 platform. Libraries were sequenced on an Illumina NovaSeq.

#### Data processing and quality controls

Gene expression, protein, hashtag, and TCR count matrices were generated using Cell Ranger, and hashtag-based demultiplexing was performed in Seurat^82,83^ . Cells were filtered based on RNA and protein quality-control metrics detailed in the immgenT Cosmology manuscript - Extended Data Note 1^29^.

#### Data integration

Datasets were integrated using totalVI^84,85^, with the 10x Genomics lane (IGT) specified as a batch covariate. The model was trained on all detected genes and proteins using a 30-dimensional latent space (default parameters).

#### Dimensionality reduction

Dimensionality reduction was performed using Minimum Distortion Embedding (MDE) implemented in the pyMDE^86^ library via the pymde.preserve_neighbors() function with default parameters. In contrast to UMAP, which is stochastic and graph-based, pyMDE preserves both local neighborhood structure and global geometry with minimal distortion. After computing the embedding on the full immgenT dataset (IGT1–96), coordinates were reused to anchor immgenT cells as a reference, enabling projection of new datasets without altering the original embedding. This enabled the construction of shared T cell reference embeddings (both all-T and lineage-specific, including Treg-specific embeddings).

#### Cell clustering and annotation

Clustering was performed in the totalVI latent space using the Louvain algorithm (Seurat FindClusters). Clustering solutions were evaluated across resolutions (0.5–4) and optimized using silhouette scores to balance over- and under-clustering. Clusters were merged or split based on consistency across samples and coherence of RNA and protein expression profiles (see immgenT Cosmology manuscript, Extended Data Note 3^29^). In total, 1,710 of 719,580 cells (0.02%) could not be confidently annotated and were labeled as “unclear” and excluded. Small provisional clusters (<1% of cells) are denoted with a “w” prefix.

Regulatory T cell clusters were identified as CD4⁺Foxp3^+^ αβ T cells based on combined RNA and protein expression of canonical markers and TFs, including *Cd3e, Trbc1, Trbc2, Trgc1, Trdc, Cd4, Cd8a, Cd8b1, Foxp3* and *Zbtb16*, as well as surface proteins CD3, TCRβ, THY1.2, CD4, CD8A, CD8B, and TCRγδ. CD4⁺Foxp3^-^ conventional αβ T cells (Tconv) formed distinct clusters.

#### CITE-seq analysis

The full panel of 128 antibodies and associated quality metrics are provided in Extended Data Table 4 and Extended Data Note 1 of the immgenT Cosmology manuscript^29^. Antibody performance was evaluated using RNA–protein correlation and protein dynamic range, stratifying antibodies into high-, intermediate-, and low-performance groups (n = 61, 41, and 26, respectively). Overall, most of the CITE-seq panel performed robustly, enabling dropout-resistant protein quantification that closely parallels flow cytometry and supports both high-dimensional analyses and conventional gating strategies. For flow cytometry–like plots, protein counts were normalized using Seurat’s LogNormalize method (log1p CP10K).

#### UMAP and clustering of individual IGT datasets

UMAP and clustering were also performed on individual IGT datasets, generating dataset-specific UMAPs used in the Rosetta2 databrowser and in some immgenT manuscripts. These used the standard Seurat functions: NormalizeData(normalization.method = "LogNormalize", scale.factor = 1e4) %>% FindVariableFeatures(selection.method = "vst", nfeatures = 2000) %>% ScaleData(features = VariableFeatures(.)) %>% RunPCA(features = VariableFeatures(.), npcs = 50) %>% FindNeighbors(dims = 1:30) %>% FindClusters(resolution = 1).

## immgenT Reference-Based Integration (T-RBI)

The full methods are described in the immgenT Cosmology manuscript^29^. Integration of query datasets using T-RBI is available at https://www.immgen.org/ImmGenT/.

Reference-based integration of external datasets (T-RBI) was performed using scVI/scANVI^85,87,88^, and pyMDE. Query datasets are first filtered for T cells using gene signature scoring, with γδ T cells identified and analyzed separately from αβ T cells. scVI models were trained jointly on reference (immgenT) and query data to learn a shared latent space, followed by scANVI to predict lineage and sub-lineage annotations with associated confidence scores. Cells with low-confidence predictions were iteratively reassigned until convergence. To quantify whether unannotated cells occupy regions of the embedding space not represented in the annotated reference (i.e., potential novel T cell states), we defined a discovery score based on a local k-nearest neighbor (kNN) distance ratio between query and reference cells. Scores >1 indicate that a cell is locally closer to other unannotated cells than to annotated cells, consistent with occupancy of under-represented or novel regions of the reference space, whereas scores ≤1 indicate embedding within previously annotated regions. The resulting latent representations were then used to project query cells onto the immgenT reference embeddings using pyMDE with anchored constraints, preserving the original atlas structure while incorporating new data.

Final outputs include lineage annotations, confidence scores, discovery scores, and coordinates in both global (all T MDE) and Treg-specific MDE embeddings, enabling direct comparison of query datasets within the immgenT framework.

## Gene Programs - Empirical Bayes matrix factorization of gene expression

To identify latent transcriptional programs, we applied empirical Bayes matrix factorization (EBMF) with a semi–non-negative matrix factorization (semi-NMF) structure to log-normalized single-cell RNA-seq expression using the flashier framework^89,90^. A detailed description of data acquisition and processing is provided in the immgenT Gene Program manuscript. Briefly, EBMF decomposes an expression matrix into a sum of low-rank components while learning sparsity, shrinkage, and noise parameters directly from the data via empirical Bayes, enabling robust identification of interpretable gene programs without requiring a priori selection of factor number or regularization strength. In the semi-NMF formulation used here, gene loadings are constrained to be non-negative, facilitating biological interpretability of gene programs, while cell loadings are unconstrained.

Gene expression was log-normalized using Seurat’s LogNormalize() with a scale factor set to the mean total UMI count per cell (mean(nCount_RNA)), yielding log1p-transformed expression values. Genes were filtered prior to model fitting to reduce technical and clonotype-driven variation: TCR variable/constant genes (e.g., Trbv/Trbj/Trbc, Trav/Traj/Trac, Trgv/Trgc, Trdv/Trdc), mitochondrial genes (^mt-), ribosomal genes (including Rpl/Rps/Mrpl/Mrps/Rsl), predicted/RIKEN/pseudogene-like features (including Gm genes, genes ending in Rik, and -ps pseudogenes), and genes not detected in any cell (row sum of counts = 0) were excluded.

Variance regularization parameters were set using a reference Poisson sampling procedure: a Poisson distribution with rate 1/n (where n is the number of cells) was sampled to estimate the standard deviation of log(x+1) and used as the residual noise scale parameter (S). EBMF components were fit using flash() (flashier^100^) with a mixture of empirical Bayes priors (ebnm_point_exponential and ebnm_point_laplace) and variance model var_type = 2, with a maximum of 200 greedily added factors (greedy_Kmax = 200). Backfitting was enabled to refine factor estimates.

## Differential Gene Expression

### Limma

Differential gene expression was performed using a pseudobulk linear modeling framework based on limma^91,92^ and edgeR. Briefly, raw RNA counts from the Seurat object were aggregated by cell cluster (annotation_level2) and experiment (IGTHT), and normalized using trimmed mean of M-values (TMM). A design matrix encoding cluster–experiment combinations was constructed, and gene-wise linear models were fitted to the normalized expression matrix using limma. Differential expression was assessed using empirical Bayes moderation of variance estimates, and statistical significance was determined with Benjamini–Hochberg correction for multiple testing. (limma_wrapper_template.sh limma_make_tmm_template.R limma_fit_template.R limma_contrasts_template.R)

### FlashierDGE

Fast differential expression was performed in the flashier^91,92^ EBMF semi-NMF framework (described in more details in the companion immgenT-GP manuscript) by leveraging the learned gene programs (factors) and cell loadings from a 200-factor model fit to log-normalized expression. For a given comparison, mean factor loadings were computed separately for group 1 and group 2, and the difference in mean loadings (Δloadings) provided differential gene-program activity. Gene-level differential expression was then reconstructed directly from the factorization as F×Δloadings, yielding per-gene log fold changes. The same framework also returned average expression estimates for both programs and genes from the group-wise mean loadings and reconstructed group means. The function is available in ZemmourLib package (FlashierDGE).

### Differential expression and gene set enrichment analysis

Differential expression (DE) between cells from each organ and the spleen baseline as well as between Treg clusters was performed using Seurat v5.3.1. Each comparison used the FindMarkers() function with the Wilcoxon rank-sum test (test.use = "wilcox"), requiring each gene to be detected in at least 10% of cells in either group (min.pct = 0.1). No log fold-change filter was applied (logfc.threshold = 0). Comparisons with empty marker tables were removed from further analysis.

## TCR Clonotype sharing and overlap analysis

Clonotype sharing across Treg clusters was assessed using matched single-cell TCR sequencing data as described in the immgenT-TCR manuscript^93^. Clonotypes were defined based on paired TCRα and TCRβ sequences with full nucleotide-level identity in the junction sequences. For each cluster, the proportion of cells with expanded clonotypes was calculated as the fraction of cells with clonotypes observed more than once in the sample. To quantify overlap between clusters, pairwise clonotype sharing was computed using the Jaccard index on sets of unique clonotypes per cluster. For visualization, a weighted network of cluster relationships was constructed, where edge weights correspond to the Jaccard index. Edges corresponding to only one shared clonotype were omitted for clarity and the network was overlaid onto the MDE embedding using cluster centroids.

## immgenT Treg flow cytometry panel design and testing

### Panel design

For each lineage, COMETSC^94^ was applied to normalized protein expression matrices together with MDE coordinates and cluster annotations, allowing up to two-gene combinations (-K = 2) to identify combinatorial markers. In combination with literature curation, extended marker combinations were evaluated for Treg cluster specificity using CITE-seq by mapping onto the Treg MDE and testing candidate strategies by flow cytometry.

### Intravascular labeling

Mice were briefly anesthetized using isoflurane and retro-orbitally injected with 3ug of CD45-PE in 200uL of PBS. Animals were euthanized after 3min as described previously^95^. Blood was drawn; colon, small intestine, spleen, and lymph nodes were harvested. Heart was perfused with 10mL of cold PBS before harvesting liver and lungs.

### Single-cell suspension from spleen, lymph nodes (inguinal, axillary, cervical, mesenteric), thymus

Immune cells were isolated using mechanical disruption followed by filtering (100uM cell strainer) and washes in PBS. An additional step for red blood cell lysis in spleen samples was performed using RBC lysis buffer (Thermofisher).

### Single-cell suspension from bone marrow

Tibias and femurs were harvested and plunged in cold 70% ethanol. Bones were rinsed in PBS, cut in half and cells were flushed out with RPMI (Gibco)-5% FBS, filtered (100uM cell strainer) and washed in PBS or RPMI-5% FBS.

### Single-cell suspension from blood

Samples were harvested by intra-cardiac puncture and mixed with heparin. 2 steps of red blood cell lysis and platelets removal by centrifugation were performed.

### Single-cell suspension from colon, small intestine

Intestinal samples were harvested and Peyer’s patches removed from the small intestine. Samples were cut in half along the intestine and in 3 pieces and incubated in RPMI containing 1 mM dithiothreitol, 20 mM EDTA, and 2% FBS at 37°C for 15min on a magnetic stirrer (500 RPM). Samples were then slightly dried to remove excess mucus, minced and dissociated in RPMI containing 1.5mg/mL collagenase II (Gibco), 0.5mg/mL Dispase II (Gibco) and 1% FBS at 37°C for 40min on a magnetic stirrer (500 RPM). Digested samples were passed through a 40uM cell strainer containing 1mL of FBS and washed in RPMI-5% FBS.

### Single-cell suspension from liver

Heart was perfused with 10mL of cold PBS. ∼2 lobes were harvested, minced and incubated in RPMI containing 1mg/mL collagenase IV (Gibco), 150ug/mL DNAse I (Sigma) at 37°C for 40min on a magnetic stirrer (500 RPM). Digested tissues were passed through a 100uM cell strained and washed in PBS. Immune cells were isolated by a Percoll (Cytiva) gradient. Briefly, cells were resuspended in 5mL of Percoll 30% and slowly transferred onto 5mL of Percoll 75% followed by a 25min centrifugation at 800g, RT without breaks. The middle ring layer was harvested, washed in PBS followed by a centrifugation at 800g, 10min, with breaks.

### Single-cell suspension from peritoneal cavity

The peritoneal cavity was infused with 5mL of cold RPMI-5% FBS, the abdomen massaged, and the volume harvested again. Cells were washed in PBS.

### Single-cell suspension from lungs

Heart was perfused with 10mL of cold PBS. Right and upper left lobes were harvested, minced and incubated in RPMI containing 1mg/mL collagenase IV (Gibco), 150ug/mL DNAse I (Sigma) at 37°C for 40min on a magnetic stirrer (500 RPM). Digested tissues were passed through a 100uM cell strained and washed in PBS. An additional step for red blood cell lysis was performed using RBC lysis buffer (Thermofisher).

### Staining

Up to 6.10^6^ cells were plated in 5mL round tubes, or P96U plate if less than 2.10^6^ cells were used. Cells were washed in PBS, and dead cells were stained using Fixable Viability Dye BUV395 (Thermofisher) for 20min in PBS at 4°C in obscurity. After a washing step in FACS buffer (PBS, 1mM EDTA, 2% FBS), cells were incubated with Fc Block (BD Biosciences) for 10min, directly centrifuged and surface antibodies were added for 20min in FACS buffer at 4°C in obscurity (**Extended Data Table 3**): CD90.1 (Thy1.1) (clone OX-7, AF488, BioLegend Cat. #202506, 1:80), FR4 (clone TH6, PE, BioLegend Cat. #125107, 1:350), TCRβ (clone H57-597, PE-Cy7, BioLegend Cat. #109222, 1:80), CD45 (clone 30-F11, BUV496, BD Biosciences Cat. #569673, 1:100), KLRG1 (clone 2F1/KLRG1, BV711, BioLegend Cat. #138427, 1:80), CD4 (clone GK1.5, BV785, BioLegend Cat. #100453, 1:80), CD304 (Neuropilin-1) (clone 3E12, PE- Dazzle594, BioLegend Cat. #145218, 1:80), CD69 (clone H1.2F3, AF700, BioLegend Cat. #104539, 1:50), CD49b (clone HMα2, APC, BioLegend Cat. #103515, 1:40), CD8a (clone 53-6.7, BUV737, BD Biosciences Cat. #612759, 1:40), CD29 (clone HMβ1-1, BV605, BD Biosciences Cat. #740365, 1:40), Sca-1 (clone D7, BV510, BioLegend Cat. #108129, 1:40), CD62L (clone MEL-14, BV650, BioLegend Cat. #104453, 1:40), CD16/CD32 (Fc block) (clone 2.4G2, purified, BD Biosciences Cat. #553141, 1:80), RORγt (clone Q31-378, BV421, BD Biosciences Cat. #562894, 1:40), FoxP3 (clone 150D, AF488, BioLegend Cat. #320012, 1:40), CD44 (clone IM7, APC, BioLegend Cat. #103012, 1:40), CD45 (clone 30-F11, PE, BioLegend Cat. #103106, 3 µg/mouse for IV labeling), CD8β (clone YTS156.7.7, BV711, BioLegend Cat. #126633, 1:80; clone H35-17.2, BV750, BD Biosciences Cat. #747505), CD44 (clone IM7, PE-Dazzle594, BioLegend Cat. #103056, 1:80). Brilliant stain buffer (Thermofisher) was added when more than 2 BV coupled markers were used.

For intracellular FoxP3 staining, cells were fixed using FoxP3 Fix/Perm buffer (eBiosciences) for 30min at 4°C in obscurity. Samples were washed in Permwash buffer 1X (eBiosciences) and resuspended with an intracellular staining cocktail for 45min at RT in obscurity: FoxP3 (clone FJK-16s, BV421, eBioscience Cat. #404-5773-82, 1:40).

Treg cells were identified as DAPI^-^ CD45^+^ TCRβ^+^ CD8α^-^ CD4^+^ Foxp3^Thy1.1/eGFP+^ using Foxp3 reporter mice (Foxp3-eGFP or Foxp3-Thy1.1) as the primary gating strategy. Where reporter mice are not available, intracellular FoxP3 staining or the CD25^hi^ CD127^low^ surface combination may be used as alternatives; however, we did not validate the panel using either of these approaches and results may vary, particularly as CD25 expression can be affected by tissue digestion protocols (e.g. dispase in gut preparations). Full antibody details including clones, providers, fluorochromes, and dilutions are provided in Extended Data Table 3, and the complete gating strategy is shown in Extended Data Fig. 3.

NovoCyte Penteon (Agilent) was used. Data were analyzed using FlowJo software version 10 (TreeStar, BD LifeSciences).

## Plotting

Plots were generated in R using ggplot2^96^, S-Plus, Seurat^83^, or the ZemmourLib R package (https://github.com/dzemmour/ZemmourLib, v0.1.3). Heatmaps were generated using pheatmap (v1.0.13) or Morpheus (Broad Institute)

## Code availability

Code is available at the following repository https://github.com/immgen/immgenT_Project/tree/main/Treg_paper

## Data availability

All raw and processed sequencing data generated in this study are available through the Gene Expression Omnibus (GEO) under accession GSE297097.

Studies mapped using T-RBI: Hanna et al.^9^, GSE196337; Muñoz-Rojas et al.^20^, GSE248440; Marin-Rodero et al.^16^, GSE234317; Yang et al.^48^, GSE188496; Dykema et al.^49^, GSE235602 ; Huang et al.^50^, GSE290617; Theune et al.^51^, GSE262405: Chi et al.^34^, GSE301231.

Additional resources are described in detail in the immgenT Cosmology manuscript^29^ (**Extended Data Note 2**).

## immgenT resource: public data browsers

Beyond the data and analyses reported in this and accompanying articles, immgenT’s goal as a public resource is served by a set of databrowsers which enable multi-faceted exploration of the data. They are accessible through the immgenT portal (https://www.immgen.org/ImmGenT/) and include:

**Rosetta2** (https://rosetta.immgen.org/) is an interactive single-cell viewer that connects RNA-seq and protein (CITE-seq) data. Cell populations can be visualized on parallel RNA and protein UMAPs, coloring reciprocally RNA- or protein-based clusters. The expression of individual genes, or of user defined gene signatures can be mapped on these representations. A cell gating interface modeled after flow cytometry serially generates 2D CITE-seq plots, on which cells can be “gated”, for projection onto UMAPs or computation of differential expression. Rosetta2 returns differentially expressed genes and proteins (as heatmaps or tables). Rosetta2 can be used to explore individual datasets or, or integrated cells grouped by lineage.

**TCR Browser** (https://rstats.immgen.org/tcrbrowser/) is a searchable interface for TCR sequences across the immgenT data. TCR repertoires can be browsed within immgenT datasets, searched for TCRs that use specific V, J or CDR3 elements, or for similarity to a user provided TCR.

**T-RBI portal** (Reference Based Integration) (https://rstats.immgen.org/immgenT_integration_analysis/) allows AI-driven mapping of external single-cel RNAseq datasets into the immgenT framework. Users upload their own data and metadata files (standard sc formats), which are then mapped and annotated within the immgenT framework. After an overnight compute, the tool returns annotation tables with integration statistics and annotation of every cell in the immgenT cluster framework, and x/y coordinates in the reference MDE space.

**Skyline browser** (https://rstats.immgen.org/Skyline/skyline.html?datagroup=immgenT%20Pseudobulk)represents, as a classic bar histogram, a pseudo-bulked overview of expression profiles of a given gene across all or selected immgenT cell clusters.

## Extended Data Figure Legends

**Extended Data Figure 1. Treg clusters characteristics.**

**(a)** Stacked bar plots showing the number of Tregs profiled in NLTs and SLOs, colored by organ.

**(b)** Stacked bar plots showing the number of Tregs profiled in male and female for each organ (experimental details in **Extended Data Table 1**).

**(c)** Bar plots showing the proportion of Treg cells profiled for each immune challenge in NLTs and SLOs, together with the associated total number of profiled Treg cells.

**(d)** Distribution of expression of the Treg-specific gene program GP68 across all T cells in the immgenT dataset.

**(e)** Treg-MDE showing the expression of *Mki67*.

**(f)** Stacked bar plots showing Treg cluster proportions in spleen, colon, skin, and thymus from healthy samples, and in spleen following NP-OVA immunization (IGT71), highlighting expansion of Treg.B. Values are shown as median ± SEM.

**(g)** Projection of the “early TCR activation” Treg cluster described by Hanna et al.^9^ onto the immgenT Treg-MDE using T-RBI (T Reference-Based Integration), shown with immgenT cluster annotations.

**(h)** Dot plot showing the mean expression (log1p(CP10K)) of effector genes across clusters. Abbreviations: AF, Afumigatus; CeD, Celiac Disease model; CRod, Crodentium infection; CT-D, CtrachomatisD infection; CT-L2, CtrachomatisL2; DSS, Dextran sulfate sodium-induced colitis; GBM-Cl, classical glioblastoma; GBM-Cl-ICB, classical GBM treated with anti-PD1 and anti- CTLA4; GBMmes, mesenchymal glioblastoma; GBM-Mes-ICB, mesenchymal GBM treated with anti-PD1 and anti-CTLA4; GBM-Pro-ICB, proneural GBM treated with anti-PD1 and anti- CTLA4; KbxN, autoimmune arthritis model; MNV-CR6, Murine norovirus CR6 strain infection; MNV-CW3, Murine norovirus CW3 strain infection; MTB, mycobacterium tuberculosis infection; PD1KO, PD-1 deficient mice; PDAC, pancreatic ductal adenocarcinoma; PDAC-ICB, PDAC treated with anti-PD1 and anti-CTLA4; PDAC-Fish, PDAC under fish oil high-fat diet; PDAC- Soy, PDAC under soybean oil high-fat diet; PP, postpartum; SA, Staphylococcus aureus (S. aureus) infection; TM, Trichuris muris helminth infection; KP, Kras-driven, p53-deficient lung adenocarcinoma model; SFB-Preg, Segmented filamentous bacteria colonization during pregnancy; Preg, pregnancy; SFB, Segmented filamentous bacteria colonization; HDM, House dust mite-induced allergic inflammation; CA, Candida albicans fungal infection; B16_ACT, B16 melanoma treated with adoptive cell therapy; Foxp3-mut, Foxp3 mutant; MF, Malassezia furfur (M. furfur) fungal infection; MCMv, murine cytomegalovirus infection; EAE, experimental autoimmune encephalomyelitis; NOD, Non-obese diabetic autoimmune diabetes model; LCMVarm, Acute LCMV Armstrong infection; B16-ACT-41BB, B16 melanoma treated with ACT, anti-PD1, and anti-4-1BB; Toxo, Toxoplasma gondii infection; VV, Vaccinia virus infection; HT- Rej, Cardiac allograft rejection; HT-Tol, Cardiac allograft tolerance; KbDb-MCMV, MCMV infection in Kb/Db-deficient mice; Qa1-KO, Qa-1-deficient mice; Qa1-MCMV, MCMV infection in Qa-1-deficient mice; LCMV-Cl13, Chronic LCMV Clone 13 infection; Lm, Listeria monocytogenes infection; MHCIIHet, MHC class II heterozygous mice; MHCIIhet_Ea16, MHCII het mice immunized with Ea16 peptide; MHCIIKO, MHC II-deficient mice; MHCIIKO_Ea16, MHCII-deficient mice immunized with Ea16 peptide; NP_OVA, Immunization with NP-OVA antigen.

Extended Data Figure 2. pTregs do not form a distinct cluster and are broadly distributed across clusters

**(a)** Treg-MDE showing Tregs sharing TCR clonotypes with Tconv cells (n = 1,462 cells), colored by cluster.

**(b)** Cluster proportion of Tregs sharing TCR clonotypes with Tconv cells (red). Box plots and black dots indicate the cluster distribution of all Tregs from the same experiment as a background distribution.

**(c)** T-RBI of pTregs from Chi et al.^34^, colored by TCR clone.

Extended Data Figure 3. Bridging flow cytometry and high-dimensional single-cell RNA sequencing using CITE-seq and the Rosetta2 website.

On the Rosetta2 website (rosetta.immgen.org), flow-like genomic analysis can be performed by gating cells based on CITE-seq protein expression and visualizing their corresponding locations on the Treg-MDE. For each panel (a–i), the left panels show Boolean gating performed on CITEseq data (gated on all Tregs in immgenT), and the right panels show the gated cells in the Treg-MDE.

**(a)** Gated CD25^hi^ IL7Rα^lo^ cells.

**(b)** Gated CD25^lo^ IL7Rα^+^ cells.

**(c)** Gated CD45RB^lo^ cells.

**(d)** Gated CD45RB^hi^ cells.

Extended Data Figure 4. Flow cytometry gating strategies for identifying Treg clusters in tissues.

**(a)** General gating strategy used to identify Treg cells.

**(b–e)** Representative gating strategies used to identify Treg.A, Treg.B, Treg.D, Treg.E and Treg.F in spleen **(b)**, liver **(c)**, colon **(d)**, and small intestine **(e)**.

Data are representative of 3–5 independent experiments.

Extended Data Figure 5. Comparing CITE-seq and flow cytometry performance for quantifying Treg clusters.

**(a)** Purity of individual Treg clusters calculated using CITE-seq (left) and flow cytometry (right).

**(b)** Scatter plot comparing the proportions of Treg.A, Treg.B, Treg.D, Treg.E, and Treg.F identified by scRNA-seq versus flow cytometry across healthy tissues. Values are shown as median ± SEM; R^2^ values and corresponding p values are indicated.

**(c)** Representative histograms of Ki-67 staining in TCRβ⁺CD4⁺CD8α⁻FoxP3⁺ Tregs from spleen, lung, small intestine, and bone marrow, highlighting proliferating Tregs. Data are representative of two independent experiments.

**(d)** Proportion of CD62L⁻ KLRG1⁻ NRP1^lo^ Sca-1⁺ Tregs (Treg.E) across thymus, spleen, colon, and small intestine. Summary statistics are shown as median ± interquartile range (n = 3). Statistical significance was assessed using a Kruskal–Wallis test with Dunn’s multiple-comparisons correction.

**(e)** Proportion of RORγt⁺ Tregs among all Tregs compared to CD62L⁻ KLRG1⁻ Sca-1⁺ NRP1^lo^ Tregs (Treg.E) in colon and small intestine (n = 3–4). Statistical significance was assessed using a Wilcoxon matched-pairs signed-rank test. Data are representative of two independent experiments.

Extended Data Figure 6. Mapping external datasets with T-RBI (T reference-based integration).

**(a,b)** T-RBI of colon Tregs from Hanna et al.^9^ **(a)** and meninges Tregs from Marin-Rodero et al.^16^

**(b)** Left, original authors’ annotations and UMAP embeddings. Right, projection of the same cells onto the immgenT Treg-MDE, shown with the original authors’ annotations (middle) and immgenT cluster annotations (right).

Extended Data Figure 7. Treg distribution across anatomical lymph nodes at baseline.

Treg-MDE plots showing cluster distributions in axillary **(a)**, inguinal **(b)**, cervical **(c)**, mediastinal **(d)**, and mesenteric **(e)** lymph nodes, colored by Treg cluster.

Extended Data Figure 8. Treg distribution during inflammation in tissues.

**(a)** Proportion of Treg clusters in healthy versus non-healthy samples. Summary statistics are shown as mean values. Statistical significance was assessed using one-way ANOVA with Sidak’s multiple-comparisons test.

**(b)** Treg-MDE plots showing cluster distributions in lung samples upon HDM challenge, flu infection, *M.tuberculosis* infection, and KP lung cancer, colored by Treg cluster.

**(c)** Treg-MDE showing expression of *Rorc* (left) and *Klrg1* (right) in healthy colon.

*HDM: House Dust Mite; KP: activation of the KrasG12D oncogene and deletion of the p53 tumor suppressor*.

Extended Data Figure 9. Treg distribution in cancer samples.

**(a)** Radar plot showing the distribution of Treg clusters in cancer across different treatments. Values are shown as median proportions.

**(b–e)** Treg-MDE with external datasets profiling tumor-infiltrating lymphocyte (TIL) Tregs mapped by T-RBI, including B16 melanoma **(b)**, orthotopic AKP tumors **(c)**, MC38 and MC38- GP colorectal cancers **(d)**, and sporadic colorectal cancer (CRC)^48–51^ **(e)**. Colored by Treg clusters.

**(f)** Radar plot showing Treg cluster distributions for the datasets shown in (b–e).

**(g)** Treg-MDE showing expression of *Ccr8* in immgenT.

Extended Data Figure 10. Treg.F characteristics.

**(a)** Left, representative gating strategy distinguishing intravascular (IV⁺), extravascular (IV⁻), and intermediate populations among CD49b⁺ CD29⁺ and CD49b⁻ Tregs in spleen (top) and liver (bottom). Right, proportion of CD49b⁺CD29⁺ and CD49b⁻ among all Tregs, with the fraction of intravascular (IV⁺) and extravascular (IV⁻) cells indicated. Values are shown as median ± SEM.

**(b)** Gene set enrichment analysis (GSEA) showing enrichment of Reactome RAP1 signaling and WP regulation of actin cytoskeleton pathways in the Treg.F transcriptional signature.

**(c)** On the Rosetta2 website, CITE-seq–based gating was used to select CD29⁺CD49b⁺ Tregs (top) and CD29⁺ CD49b⁻ Tregs (bottom), which were then projected onto the Treg-MDE. This analysisshows that CD29⁺ Tregs comprise two distinct populations, CD49b⁺ and CD49b⁻, indicating that CD29 expression alone is insufficient to define Treg.F.

**(d,e)** Proportion **(d)** and absolute numbers **(e)** of CD62L^-^ KLRG1^-^ CD49b^+^ CD29^+^ Tregs in the spleen and liver of WT and Foxp3^Cre^ Klf2^fl/fl^ mice (n = 7 across four experiments). Horizontal bars indicate the median, and statistical significance was assessed using a two-sided Mann–Whitney *U* test.

Extended Data Figure 11. Treg cells in Aβ^+/o^ and Aβ^o/o^ mice

**(a)** Treg-MDE showing the Treg cluster distributions in Aβ^+/o^ and Aβ^o/o^ mice from IGT91/92.

**(a)** Treg Vα and Vβ usage in Aβ^+/o^ and Aβ^o/o^ mice from IGT91/92.

**Extended Data Table 1. Metadata information about the immgenT samples. Extended Data Table 2. Treg cluster gene signatures.**

**Extended Data Table 3. immgenT Treg flow cytometry panel.**

**Extended Data Table 4. Correspondence between immgenT clusters and common annotations**

