## Extended Data Figures for "A Reference Landscape of Regulatory T Cell States in Mice"

Extended Data Figure 1

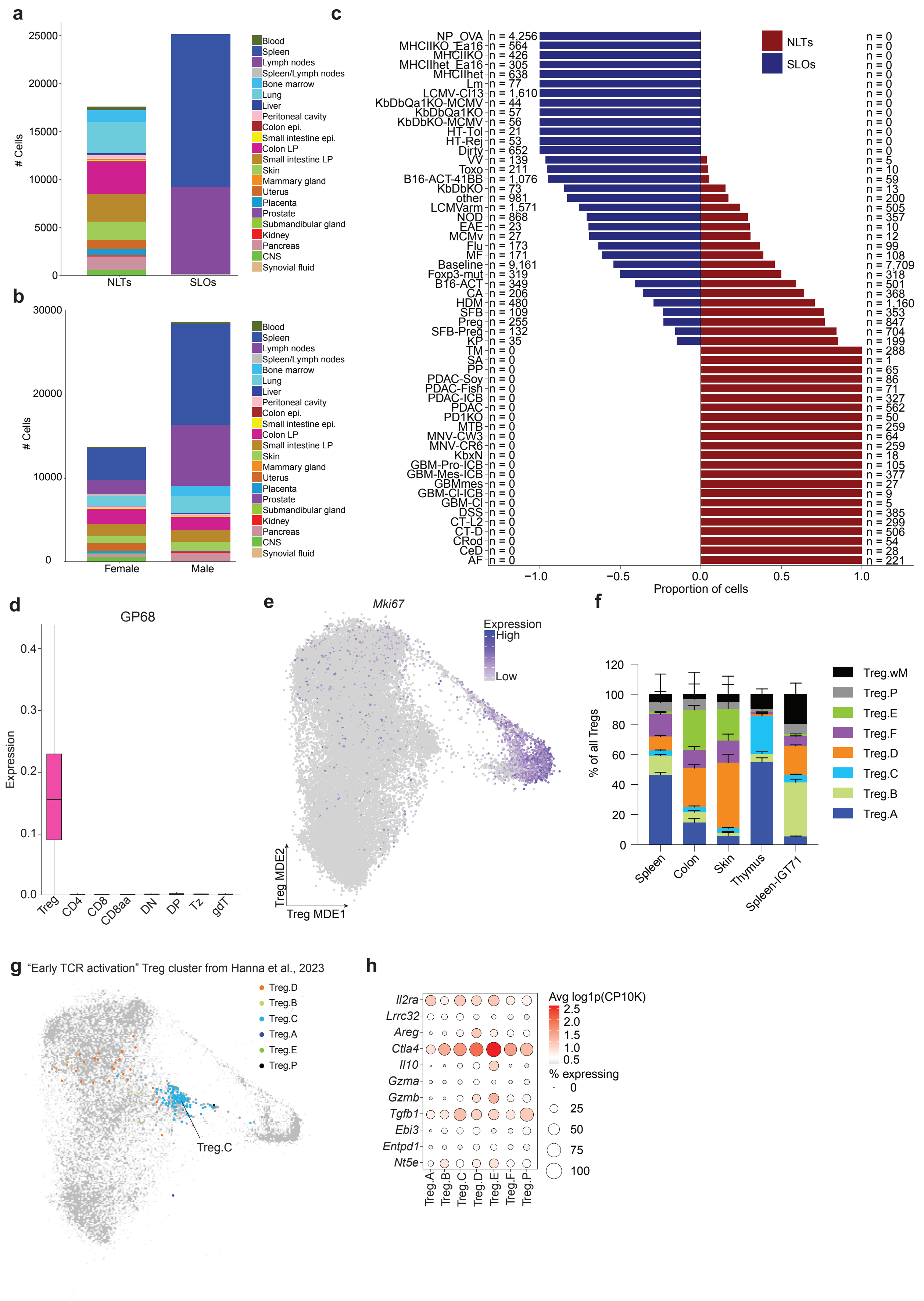

Extended Data Figure 2

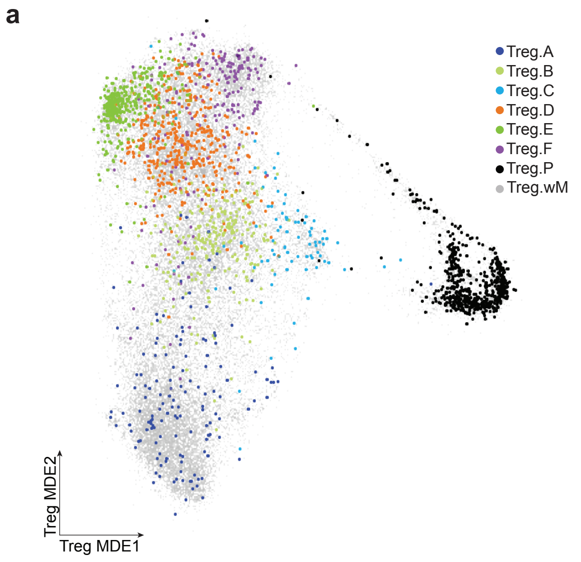


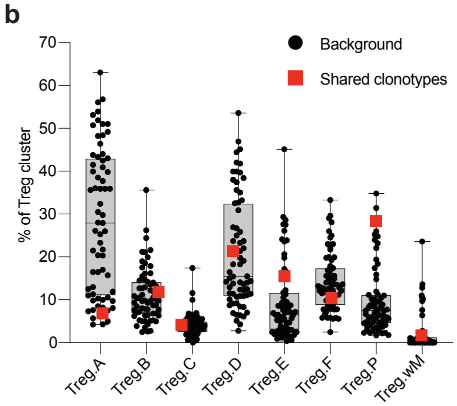


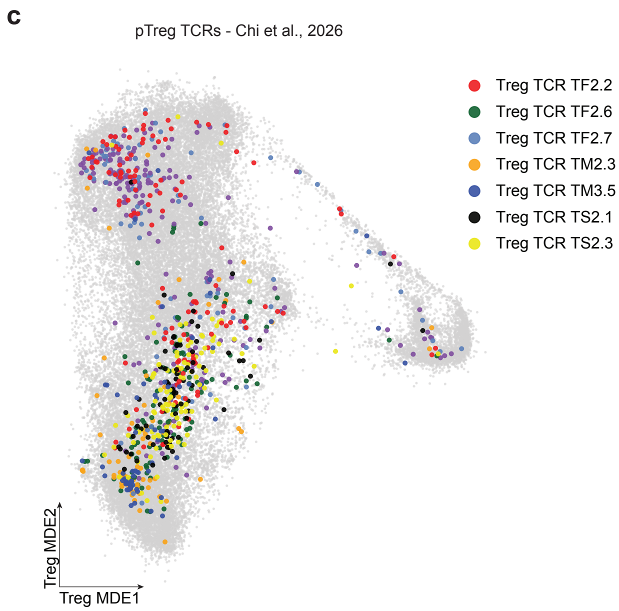

Extended Data Figure 3

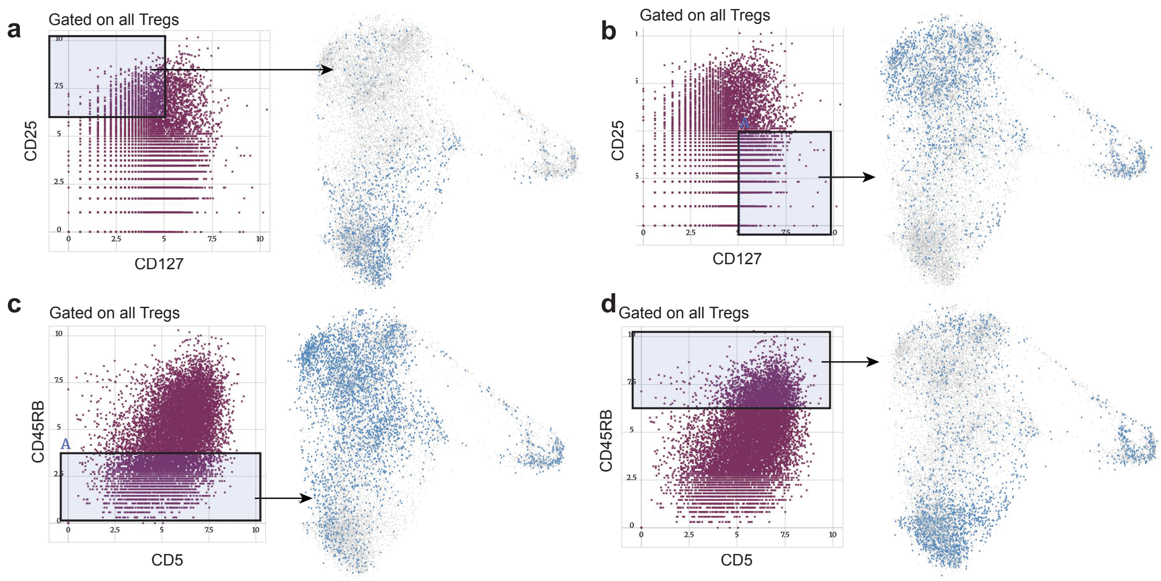

Extended Data Figure 4

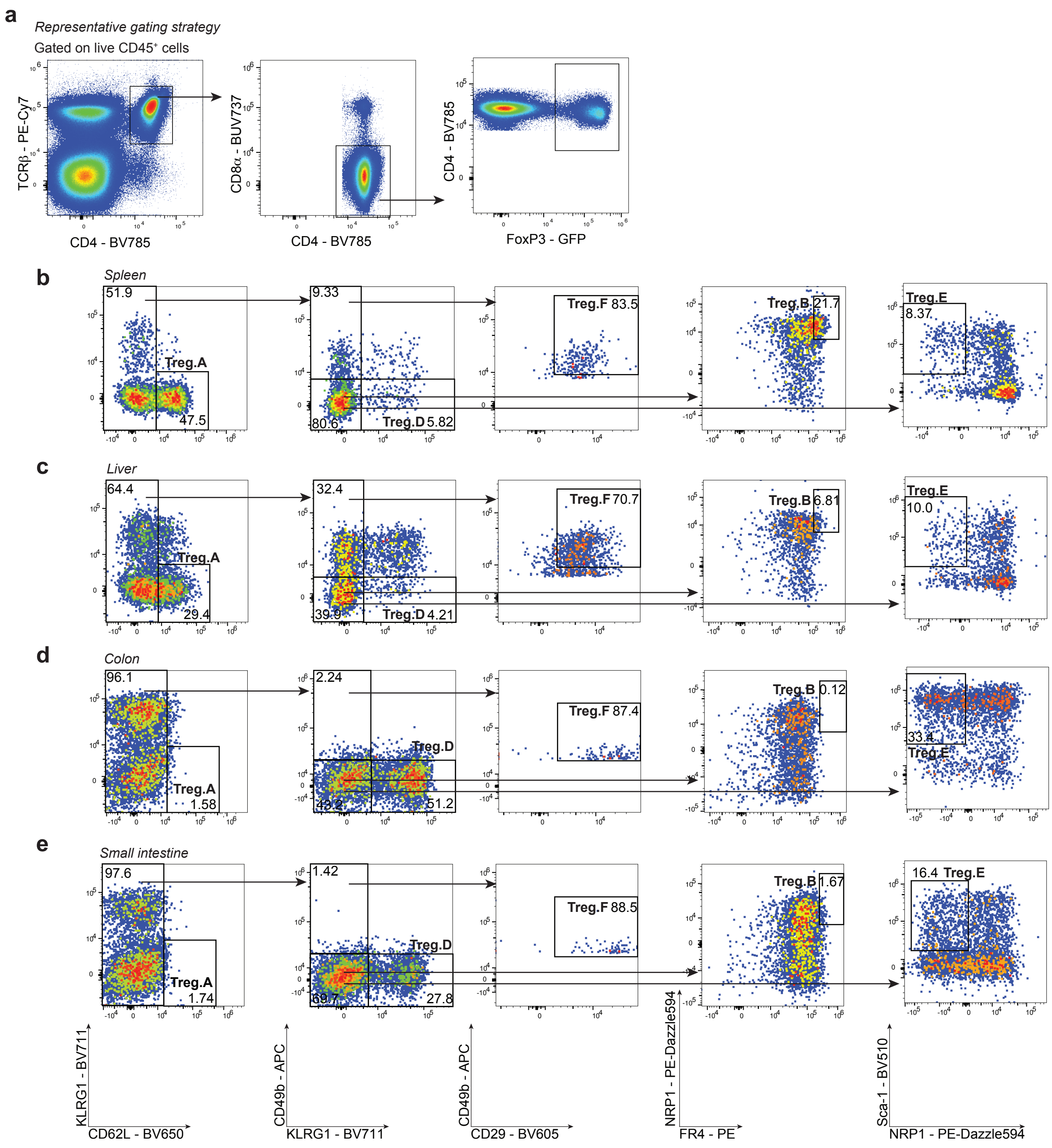

### Extended Data Figure 5

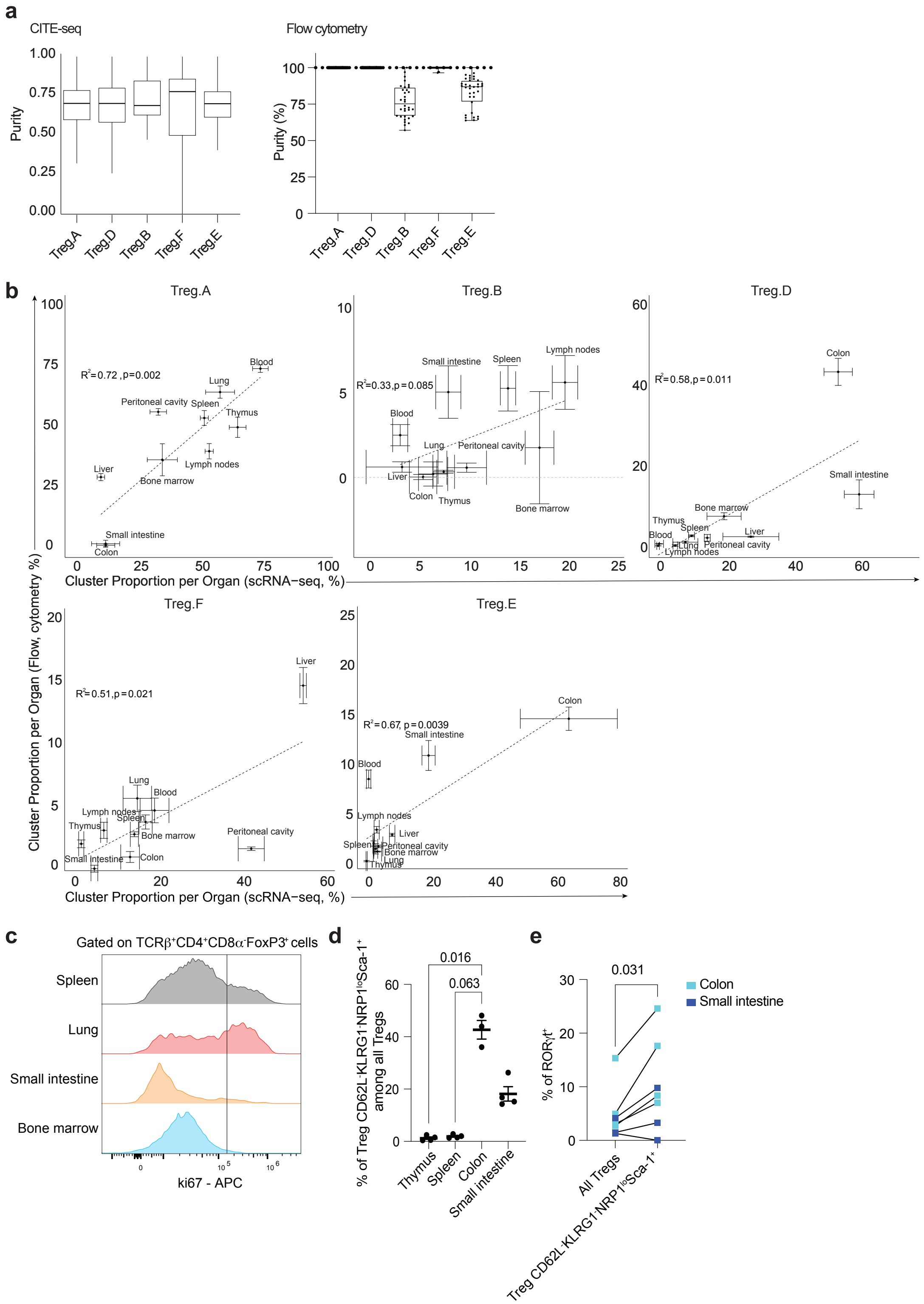

Extended Data Figure 6

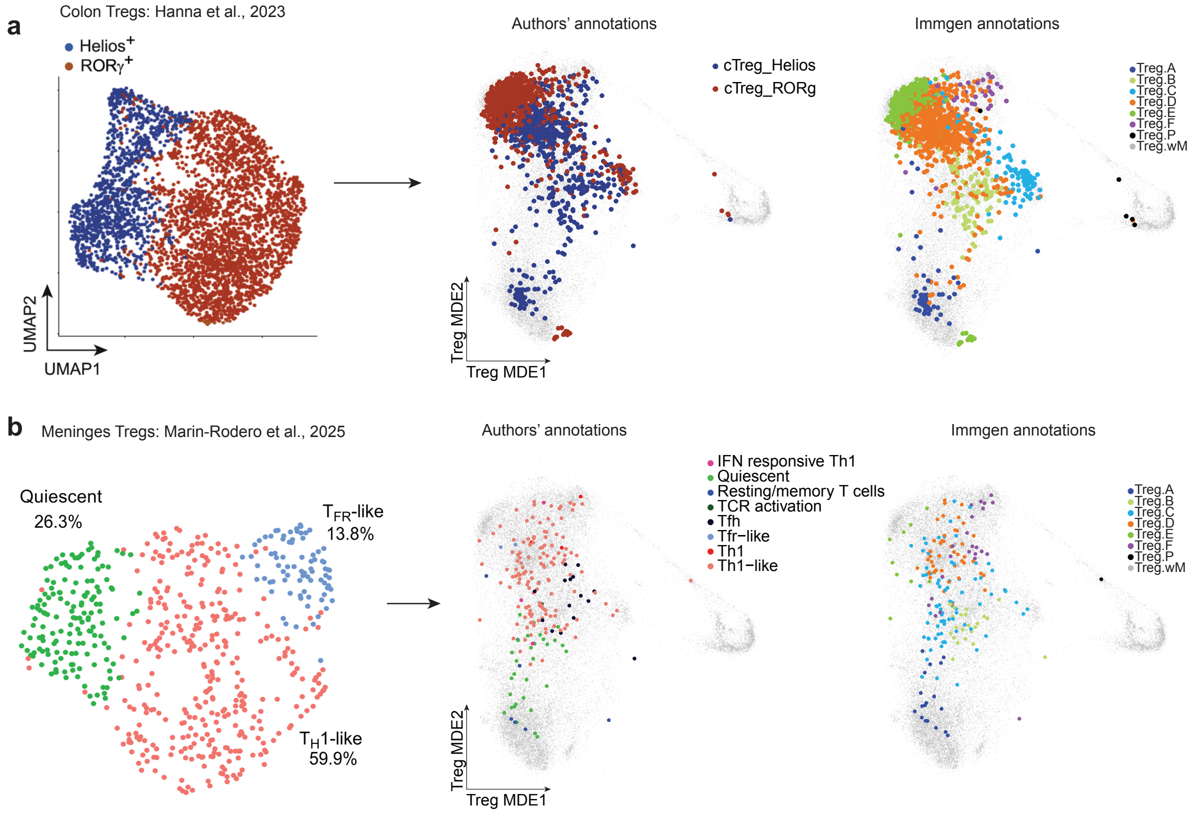

Extended Data Figure 7

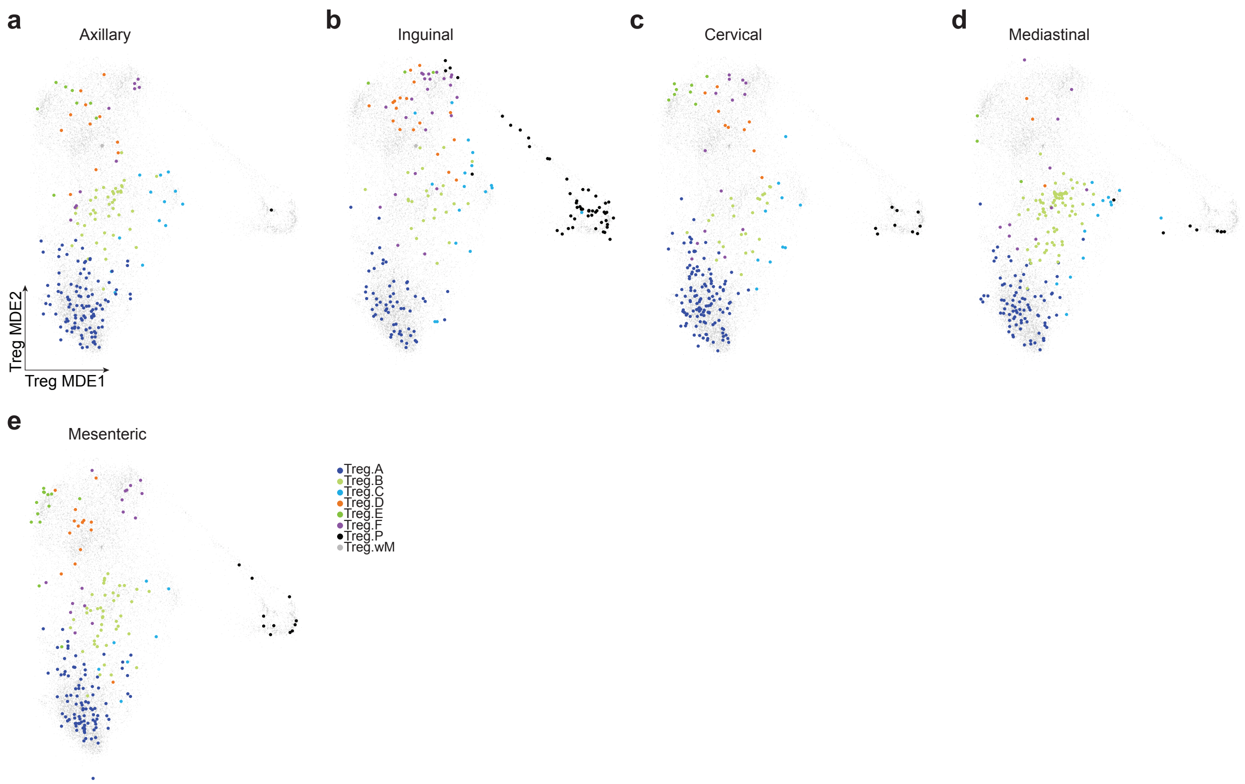

Extended Data Figure 8

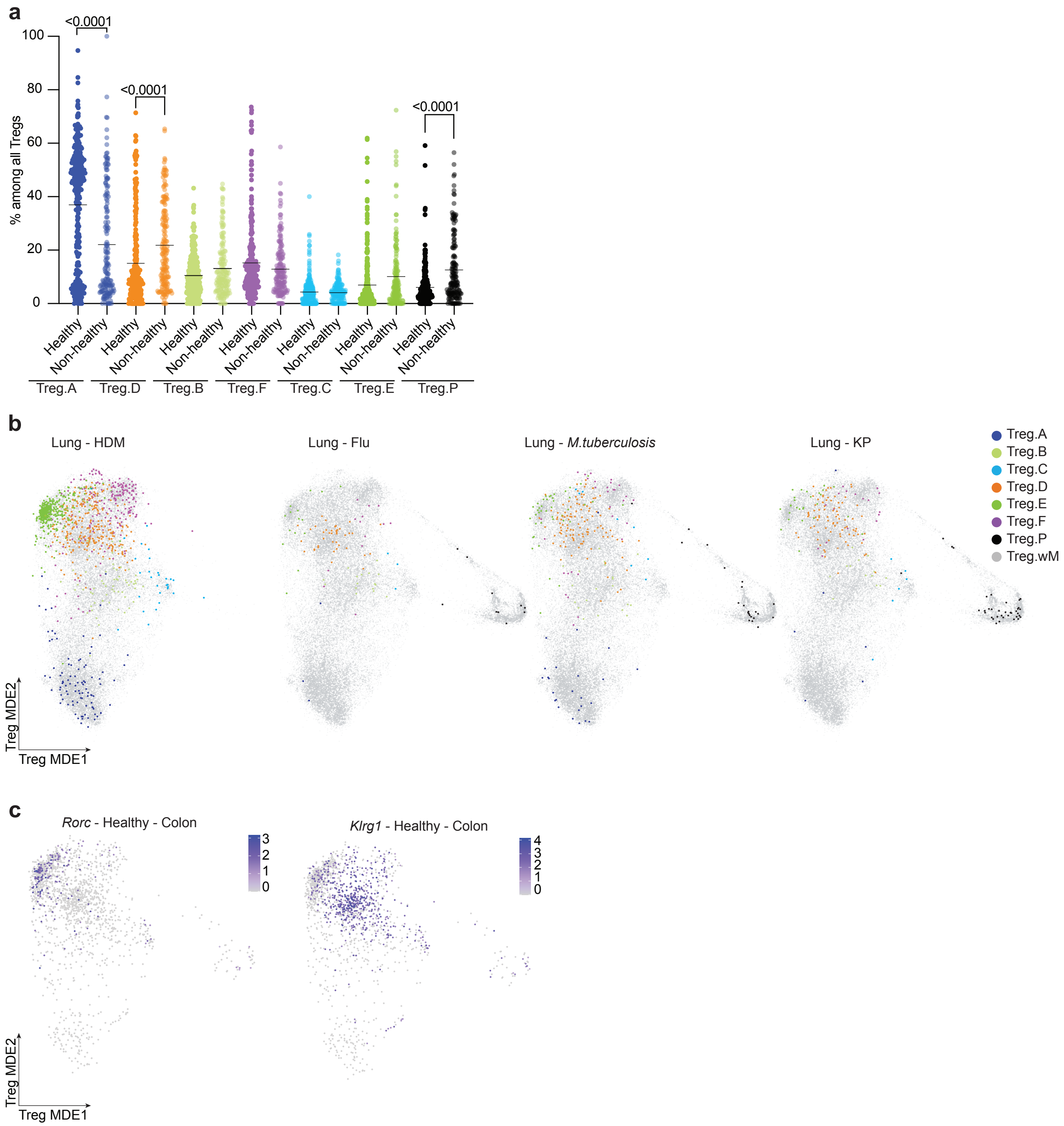

Extended Data Figure 9

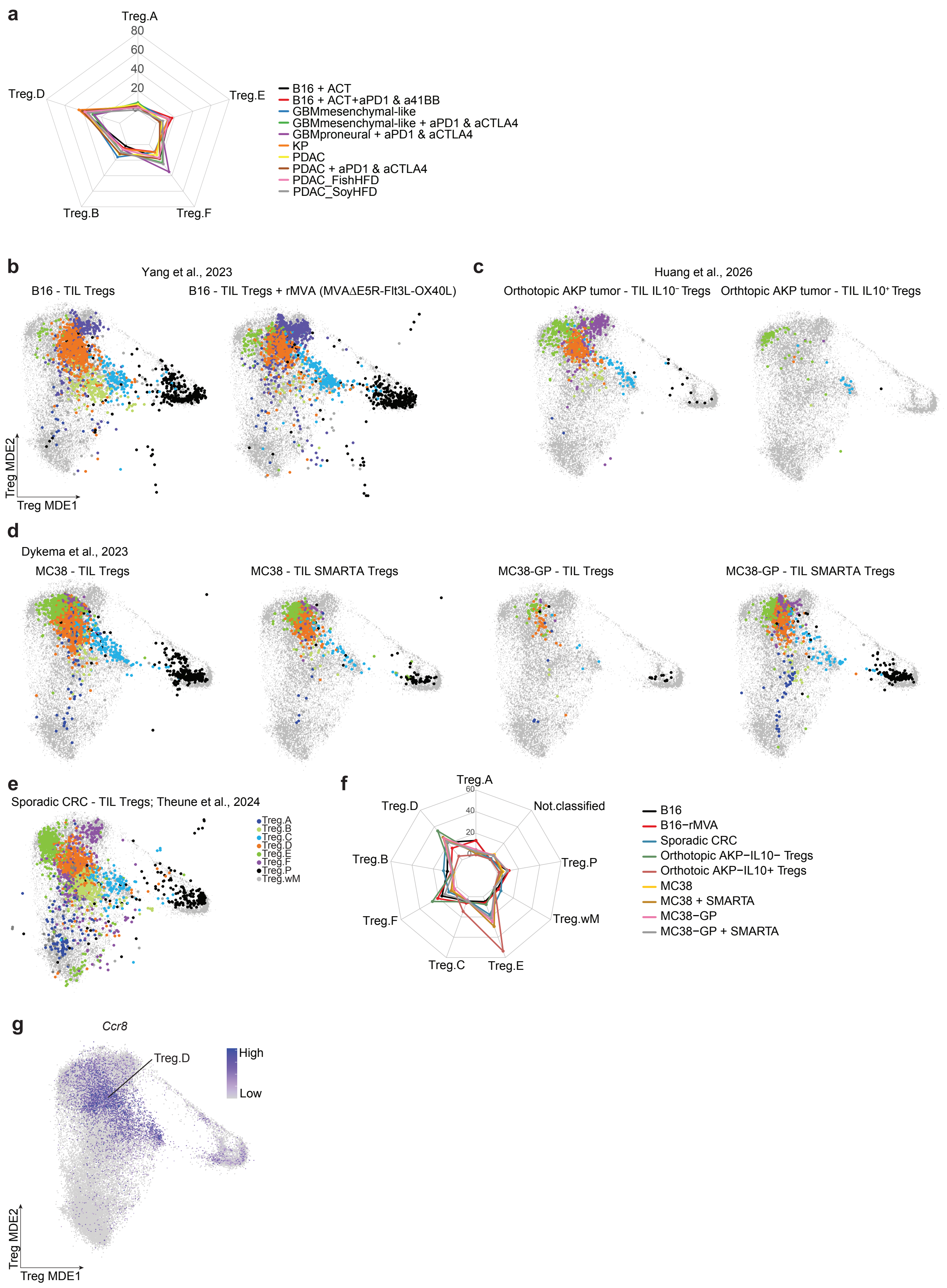

### Extended Data Figure 10

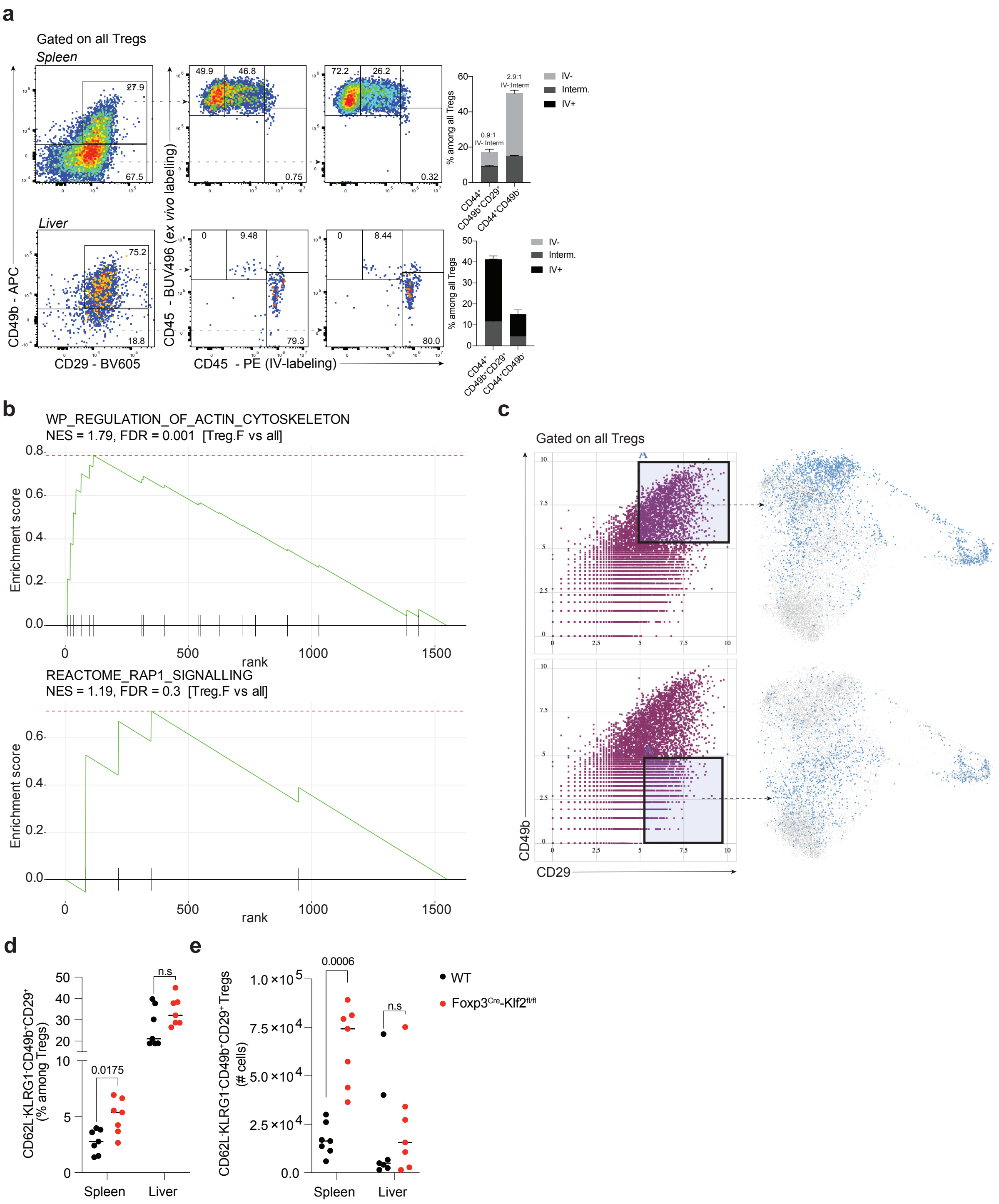

Extended Data Fig 11

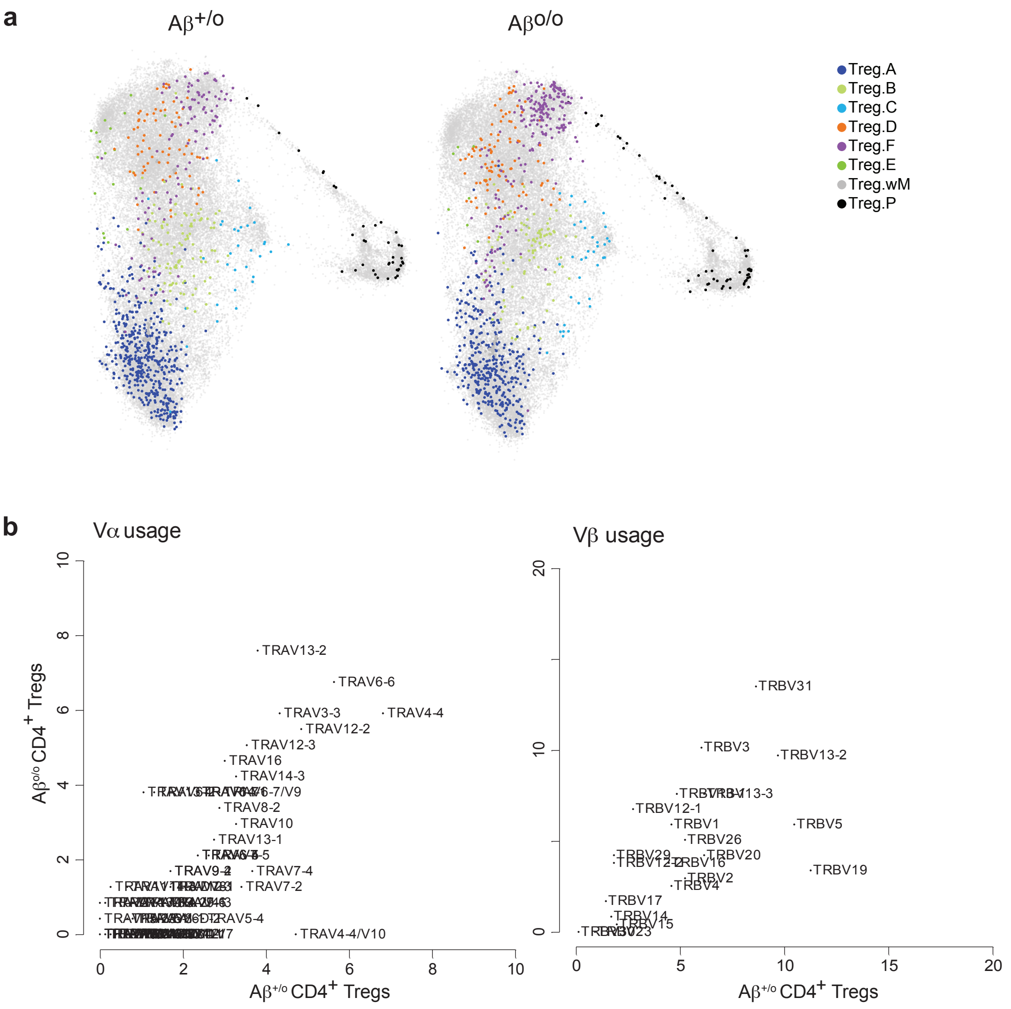
